# A thalamic perspective of (un)consciousness in pharmacological and pathological states in humans

**DOI:** 10.1101/2024.05.30.596600

**Authors:** Dorottya Szocs, Dian Lyu, Andrea I. Luppi, Peter Coppola, Rebecca Woodrow, Guy B. Williams, Judith Allanson, John D. Pickard, Adrian M. Owen, Lorina Naci, David K. Menon, Emmanuel A. Stamatakis

## Abstract

Currently, there is substantial ongoing discussion around the functional role of the thalamus in consciousness. What is missing in the literature, however, is a systematic investigation of the relevance of specific thalamic nuclei in pharmacologically and pathologically altered states of consciousness in humans. Using functional neuroimaging in both healthy anaesthetised volunteers and patients with disorders of consciousness (DOC), we sought to identify which specific thalamic subregions in both cohorts may be differentially significant for loss of consciousness. Our findings revealed that the pulvinar (Pu) and ventral-latero-ventral (VLV) nuclei, in anaesthesia, and the VLV, in DOC, had distinct functional connectivity patterns related to the default mode and somatomotor networks. Remarkably, among all nuclei, the Pu was found to have the strongest functional connectivity change with anaesthetic-induced loss of consciousness, while in DOC patients, we found the VLV revealed the strongest connectivity change in comparison with healthy controls. Furthermore, we provide evidence that this neural connectivity biomarker in patients also mirrors the changes observed at the behavioural level, which could have clinical implications for targeted deep brain stimulation in therapy for DOC.

## Introduction

The quest to identify the ‘neural correlates of consciousness’ has both scientific and clinical relevance, given the pressing challenge of detecting covert consciousness in uncommunicative individuals with disorders of consciousness.^1^ We are currently still lacking a definitive understanding of which neural circuits — common or differential — are responsible for altered states of consciousness and how these work.^2^ There is substantial evidence that functional neuroimaging can not only detect covert awareness in patients who appear entirely unresponsive at the bed side, but can predict whether a patient may recover.^3^

Recent studies have shown, the thalamus is involved in the modulation of arousal states.^4,5^ This complex subcortical structure has received particular attention as it is situated, both anatomically and functionally, at the convergence of the hypothalamus, telencephalon, and brainstem, and most accounts of its functional role propose that it coordinates dynamic interactions between the cortex, basal ganglia, and cerebellum, which form the basis of our conscious experience.^4,6^ Even more so, at a macroscopic level, it plays a crucial role in systems-level dynamics by integrating multimodal information across large-scale cortical networks.^7^ Furthermore, specific thalamic nuclei mediate higher-order cognitive functions,^8^ such as attention^9,10^ and awareness.^11,12^ Consciousness has been investigated extensively from the perspective of large-scale cortical networks.^13^ Among these networks, the default mode network (DMN) is well-known to be largely linked to conscious states.^14,13,15^ It is a widely distributed network consisting of medial frontal and posterior medial parietal cortices, angular gyrus, and hippocampus, of which the posteromedial cortex is a major connectivity hub^16^ and has been increasingly associated with mediating internal (self-related) and external (environmental) information processing.^17,18^ The posteromedial area has been demonstrated to play a key role in altered consciousness in epileptic patients^19,20,21^ and in patients with disorders of consciousness.^22,23,24^

A growing body of animal literature has demonstrated the role of distinct thalamic nuclei in pharmacologically induced unconsciousness, specifically demonstrating that central thalamic stimulation induced arousal in anaesthetised macaques.^25,11^ Equally, human studies have highlighted thalamic interactions with the default mode network (DMN) in anaesthesia and disorders of consciousness.^26,22^ However, only few human studies have differentiated the cytoarchitecturally and functionally distinct thalamic nuclei, to elucidate their respective relevance in altered states of consciousness.^27,28,29^ There have been few systematic studies of the neuropathology of the vegetative state, particularly as affecting the thalamus. Importantly, all but one of 24 VS patients who survived for more than three months after traumatic brain injury (TBI) had thalamic damage (96%) that was mainly attributable to diffuse axonal injury, compounded in some cases by secondary insults including hypoxia and raised intracranial pressure.^30^ Moreover, neuronal loss occurred in the mediodorsal parvocellularis, rostral center medial, central lateral and paracentral nuclei in moderately disabled patients; and from the mediodorsal magnocellularis, caudal centre medial, rhomboid, and parafascicular nuclei in severely disabled patients; and all of the above and the centre median nucleus in VS patients.^31^Additionally, neuronal loss occurred primarily from cognitive and executive function nuclei, a lesser loss from somatosensory nuclei and the least loss from limbic motor nuclei.^31^

What is missing, however, is a thorough investigation of the differential role of individual thalamic nuclei with relevance to pathological impairment in consciousness following brain injury. This is critical given the current active debate around the functional role of the thalamus in consciousness.^4,32^ Further, such understanding is crucial from a clinical standpoint, since deep brain stimulation (DBS) of the central-lateral (CL) nuclei has been found to improve behavioural outcomes in DOC patients.^33,34^ Thalamic stimulation has been implemented in other brain disorders, notably obsessive-compulsive disorder,^35^ Parkinson’s disease, essential tremor, and dystonia.^36^ More recently, a stimulation paradigm of both the anterior and medial pulvinar thalamic nuclei has been efficacious for treating drug-resistant epilepsy.^37,38^ Even more so, with advances in MRI spatial resolution and improvements in thalamic parcellation atlases,^39,40,41,42,43,44,45^ we are witnessing innovative attempts to functionally disentangle the thalamic ensemble. This has led to more investigations of the role that individual thalamic nuclei play in the context of arousal, most notably in sleep^28^ and propofol anaesthesia.^27^

To our knowledge, this study is the first systematic investigation of distinct thalamic nuclei i.e. 7 major clusters — pulvinar (Pu), anterior (Ant), medio-dorsal (MD), ventral-latero-dorsal (VLD), central-lateral, lateral-posterior, medial-pulvinar (CL-LP-MPu), ventral-anterior (VA), and ventral-latero-ventral (VLV) — and their functional relationship to cortical and subcortical areas, using functional neuroimaging datasets from cohorts of (i) healthy anaesthetised volunteers (n=16) i.e. pharmacologically-induced; and (ii) patients with disorders of consciousness (n=22) i.e. pathological, given that anaesthesia has been previously demonstrated to be an experimental model for DOC studies.^46,47,48^ Crucially, we sought to pinpoint which thalamic nucleus in both cohorts is most strongly associated with loss of consciousness, which may, in turn, shed light upon the neural mechanisms of conscious processing, and more crucially, indicate potential areas for targeted brain stimulation therapy in disorders of consciousness.

## Methods

### Data acquisition

#### Anaesthesia London Ontario Dataset

The original study^49^ acquired the anaesthesia dataset between May and November 2014 at the Robarts Research Institute in London, Ontario (Canada), approved by the Western University Ethics board, and have been previously published before.^47,50,22,52^ 19 healthy (13 males; 18–40 years), right-handed, English speakers with no reported neurological conditions signed an informed-consent sheet. Due to equipment malfunction or impairments with the anaesthetic procedure, 3 participants were excluded, thus, 16 participants were included in this study.^49^

Resting-state fMRI data were acquired at different propofol levels: no sedation (Awake), Deep anaesthesia (corresponding to Ramsay score of 5) and during post-anaesthetic recovery. For each condition fMRI acquisition began after two anaesthesiologists and one anaesthesia nurse independently assessed Ramsay level in the scanning room. The anaesthesiologists and the anaesthesia nurse could not be blinded to experimental condition, since part of their role involved determining the participants’ level of anaesthesia. The Ramsay score is designed for critical care patients, and therefore participants did not receive a score during the Awake condition before propofol administration; rather, they were required to be fully awake, alert and communicating appropriately.^22,49^

Propofol was administered intravenously using a Baxter AS50 (Singapore); stepwise increments were applied via a computer-controlled infusion pump until Ramsay level 5 was reached (i.e. no responsiveness to visual or verbal incitements stimulation). If necessary, further manual adjustments were made to reach target concentrations of propofol which were predicted and maintained stable by a pharmacokinetic simulation software TIVA trainer (European Society for Intravenous Anaesthesia, eurosiva.eu). Blood concentration levels were measured following the Marsh 3-compartment model. The initial propofol concentration target was 0.6 *𝜇*g/ml, and step-wise increments of 0.3 *𝜇*g/ml were applied after which Ramsay score was assessed. This procedure was repeated until participants stopped responding verbally and were rousable only by physical stimulation at which point data collection would begin. Oxygen titration ensured SpO2 above 96%. The mean estimated effect-site propofol concentration was 2.48 (1.82–3.14) µg mL^−1^, and the mean estimated plasma propofol concentration was 2.68 (1.92–3.44) µg mL^−1^. Mean total mass of propofol administered was 486.58 (373.30–599.86) mg.^22,49^ 8 min of RS-fMRI data was acquired. A 3-tesla Siemens Trio scanner was used to acquire 256 functional volumes (Echo-planar images [EPI]). Scanning parameters were: slices = 33, 25% inter-slice gap resolution 3 mm isotropic; TR = 2000 ms; TE = 30 ms; flip-angle = 75°; matrix = 64 × 64. Order-of-acquisition was bottom-up interleaved. The anatomical high-resolution T1 weighted images (32-channel coil, 1 mm isotropic voxels) were acquired using a 3D MPRAGE sequence with TA = 5 mins, TE = 4.25 ms, matrix = 240 × 256, 9 degrees flip angle. ^22,49^

#### Disorders of Consciousness Dataset

MRI data for 24 DOC patients were collected between January 2010 and July 2015 in the Wolfson Brain Imaging Centre in Addenbrookes Cambridge, UK. For the present study, these patients were selected out of a bigger dataset (*n* = 71) due to their relatively intact neuroanatomy.^46,47,48,50^ These patients were admitted to the research ward and scanned at the Wolfson Brain Imaging Centre, Addenbrookes Hospital (Cambridge, UK). Written informed assent was obtained from the referring clinical teams and a family member or other relevant close contact. Following a full neurological examination and daily behavioural observations using the Coma Recovery Scale-Revised (CRS-R), participants were allocated a diagnosis of unresponsive wakefulness syndrome (UWS) or minimally conscious state (MCS). Patients whose behavioural responses were not indicative of awareness at any time, were classified as UWS. In contrast, patients were classified as being in a minimally conscious state (MCS) if they demonstrated behavioural evidence of simple automatic motor reactions (e.g., scratching, pulling the bed sheet), visual fixation and pursuit, or localisation to noxious stimulation). ^46,47,48^ MCS patients were further classified depending on the presence (MCS+) or absence (MCS-) of language function.^51^ Detailed demographics are presented in Table 1. The collection of this dataset received ethical approval from the National Research Ethics Service. ^46,47,48^

**Table 1.** Demographics for DOC patients

| Patient | Gender | Age | Aetiology | Months Post-Injury | Diagnosis | CRS | Tennis Task |
| --- | --- | --- | --- | --- | --- | --- | --- |
| 1 | M | 21 | TBI | 45 | MCS+ | 11 | positive |
| 2 | M | 57 | TBI | 14 | MCS- | 12 | negative |
| 3 | M | 47 | TBI | 4 | MCS+ | 10 | negative |
| 4 | M | 36 | TBI | 34 | UWS | 8 | negative |
| 5 | M | 17 | Anoxic | 46 | UWS | 11 | negative |
| 6 | F | 38 | Anoxic | 9 | MCS- | 10 | negative |
| 7 | F | 38 | TBI | 13 | MCS+ | 11 | positive |
| 8 | M | 29 | TBI | 68 | MCS+ | 10 | positive |
| 9 | M | 23 | TBI | 4 | MCS+ | 7 | positive |
| 10 | F | 70 | Cerebral bleed | 11 | MCS+ | 9 | negative |
| 11 | F | 30 | Anoxic | 6 | MCS- | 9 | positive |
| 12 | F | 34 | Anoxic | 6 | UWS | 8 | negative |
| 13 | M | 22 | Anoxic | 5 | UWS | 7 | negative |
| 14 | M | 37 | Anoxic | 14 | UWS | 7 | negative |
| 15 | F | 62 | Anoxic | 7 | UWS | 7 | negative |
| 16 | M | 46 | Anoxic | 10 | UWS | 5 | negative |
| 17 | M | 21 | TBI | 7 | MCS+ | 11 | negative |
| 18 | M | 67 | TBI | 14 | MCS- | 11 | positive |
| 19 | F | 55 | Hypoxic | Unknown | UWS | 12 | negative |
| 20 | M | 28 | TBI | Unknown | MCS+ | 8 | positive |
| 21 | M | 22 | TBI | Unknown | MCS+ | 10 | negative |
| 22 | F | 28 | ADEM | Unknown | UWS | 6 | negative |

Patients were further stratified into two groups based on their ability to perform volitional tasks (mental imagery) in the scanner which have been previously used to assess the presence of covert consciousness in DOC patients.^46,54,55,56^ Patients were instructed to perform two mental imagery tasks. The first task involved motor imagery (“tennis task”), where each patient was asked to imagine being on a tennis court swinging their arm to hit the ball back and forth with an imagined opponent. The second task involved spatial imagery (“navigation task”), where the patient was required to imagine walking around rooms of their house, or streets of a familiar city, and to visualise what they would see if they were in those locations. Each task comprised five cycles of alternating imagery and rest blocks and lasted 30 seconds. The two types of blocks were cued with the spoken word “tennis” or “navigation”, respectively, whereas the resting blocks were cued with the word “relax”, corresponding to instructions for the patient to just stay still and keep their eyes closed. Univariate fMRI analysis was conducted on all patients for both tasks and the analysis was conducted in FSL (https://fsl.fmrib.ox.ac.uk/fsl/fslwiki/); for each functional scan, a general linear model (GLM) with alternating blocks of rest and active imagery was calculated.^53^ Results were considered significant at a cluster level of z > 2.3 (corrected *p* < 0.05 at the cluster level).^46,53^ Some patients showed significantly greater brain activation during either of the “tennis task” (supplementary motor area) or “navigation task” (parahippocampal place area) than rest, which was used as evidence for task-responsiveness.^46,53^ These patients, thus, were categorised (Table 1) as “fMRI+ positive task responders” (labelled ‘positive’), whereas the rest were “fMRI-negative non-responders” (labelled ‘negative’).

Resting-state fMRI was acquired for 10 min (300 volumes, TR = 2 s) using a siemens TRIO 3T scanner. The functional images were acquired using an echo planar sequence. Parameters include: 3 × 3 × 3.75 mmm resolution, TR/TE = 2000 ms/30 ms, 78° FA. Anatomical images T1-weighted images were acquired using a repetition time of 2300 ms, TE = 2.47 ms, 150 slices with a cubic resolution of 1 mm. In the present study, patients were systematically excluded from the final analysis based on the following criteria: (1) presence of large focal brain damage (greater than one third of a hemisphere) and thalamic damage as assessed by a neuroanatomical expert; (2) excessive head motion during resting-state scanning (greater than 3 mm in translation and/or 3 degrees in rotation); and (3) segmentation and normalization failure during pre-processing^22,47,48,50^ (Table 1).

#### Data pre-processing

All functional and structural images were pre-processed using SPM12 (https://www.fil.ion.ucl.ac.uk/spm/software/spm12/). The first 5 functional scans were removed to reach scanner equilibrium/steady-state magnetisation. Slice-timing correction was performed on the fMRI volumes, followed by realignment to the mean functional volume, which produced realignment parameters that were included in the first-level statistical models. Using the mean functional image, direct spatial normalisation to an EPI-template was performed using the function ‘old norm’ in SPM due to reduced variability across subjects compared to other approaches in previous studies.^57^ Participant’s high-resolution structural images were co-registered to the mean functional image or mean EPI (produced from the realignment process) and segmented into grey matter, white matter, and cerebrospinal fluid masks, and finally spatially normalised to the MNI-152 template.^58^ Visual inspection or quality control of the images was conducted for normalisation to the standard space, which was essential for the DOC dataset due to the potential effect of lesions on spatial transformations. To avoid bias in the results, at this stage, 1 subject was excluded due to the half of the brain missing from the volumes, and 1 subject was excluded due to complete failure of normalisation. Denoising was then performed in the Matlab-SPM-based software CONN (https://web.conn-toolbox.org), and functional images were smoothed with a 6mm FWHM Gaussian kernel. Movement parameters and their first temporal derivative were included as a first-level covariate to remove motion-related noise. The aCompCorr algorithm regressed out CSF, white-matter, and motion-related signals from the time-series (using the first 5 principal components). This method has been shown to perform well in removing movement, respiratory and cardiac artefacts,^59^ especially on DOC patients,^60^ and has been previously used for this type of data.^47,61,22^ The ART quality-assurance/motion-artefact rejection toolbox (https://www.nitrc.org/projects/artefact_detect), as implemented in CONN, was also used to further remove motion-related artifacts in the time-series data. This method involves regressing out the effect of outlier scans (movement *>* 0.09 mm) in a first-level analysis which is suggested to further reduce focal effects of movement not accounted for by aCompCorr algorithm.^59,62^ Linear detrending and 0.008–0.009 Hz band-pass filter was applied.

#### Thalamic parcellation atlas

We adopted seed-based functional connectivity (FC) analysis to establish which brain regions, specific thalamic nuclei functionally interact with. In seed-to-voxel FC analysis, the desired seeds — in this case, thalamic nuclei — need to be defined in regions-of-interest (ROIs).^63^ While MRI provides an enhanced depiction of soft biological tissue and allows delineation and characterisation of distinct structures, the intrinsic contrast provided by T1- and T2-weighted MRI is too low to be able to differentiate between distinct nuclei, especially given the relatively small size of the thalamus (8cm^3^ per hemisphere), which highlights the need for developing thalamic parcellation methods. Currently, there are a limited number of digital atlases of the thalamus, with minimal cross-correspondence between them.^40^ Recently, a probabilistic atlas of the anatomical subdivisions of the thalamus built on 70 healthy subjects from the Human Connectome Project has been developed based on diffusion-weighted MRI.^41^ DW-MRI is the only non-invasive imaging technique able to depict white matter fibre characteristics within each thalamic nucleus with respect to cortical projections. In this atlas^41^ the thalamus is segmented into seven regions closely matching the anatomical subparts, where six clusters correspond to histologically defined larger thalamic nuclei and the seventh cluster is a conglomerate of three nuclei. We used a thalamic parcellation atlas based on DW-MRI, where the thalamus is segmented into seven regions of interest (ROIs) closely matching the anatomical subparts.^41^ We chose to use this parcellation as it is a good trade-off between the spatial position of thalamic voxels and their local diffusion properties and has demonstrated robust similarity to the thalamic anatomy presented in Morel’s stereotactic atlas.^41,64,65^ Atlas standard space of the thalamus probabilistic masks was transformed into the preprocessed standard space using the nearest neighbour interpolation, in order to avoid overlapping of the thalamic masks and impact of results.^66^ Thalamic masks used, therefore, correspond the following bilateral nuclei: pulvinar (Pu), anterior (Ant), medio-dorsal (MD), ventral-latero-dorsal (VLD), central-lateral, lateral-posterior, medial-pulvinar group (CL-LP-MPu), ventral-anterior (VA), and ventral-latero-ventral (VLV) (See Supplementary Fig. 1).

#### Seed-to-voxel FC and intrinsic connectivity networks (ICNs)

We analysed the fMRI data of N=16 healthy controls under deep sedation (mean plasma propofol concentration of 2.68𝜇g/mL) and N=22 disorders of consciousness (DOC) patients. Thalamic masks corresponded to the following nuclei (bilaterally): pulvinar (Pu), anterior (Ant), medio-dorsal (MD), ventral-latero-dorsal (VLD), central-lateral, lateral-posterior, medial-pulvinar (CL-LP-MPu), ventral-anterior (VA), and ventral-latero-ventral (VLV).

Functional connectivity was calculated using CONN (https://web.conn-toolbox.org) in the form of seed-to-voxel analyses from 7 seeded thalamic nuclei (Supplementary Fig. 1) aiming to investigate thalamic nuclei interactions with the rest of the brain. Temporal correlations for the thalamic seeds were computed for all other voxels in the brain using a general linear model (GLM). For the anaesthesia dataset where the experimental design is within-subject, functional connectivity analyses were first conducted individually, then the individual seed-to-voxel parameter estimate images were entered into group-level analyses. Specifically, paired-sample *t* tests were used for differences in the participants among the conditions of deep sedation vs. awake and recovery vs. deep sedation. The cortical SPM-*t* maps from the individuals, both unthresholded and thresholded, were grouped together with a one-sample t-test for the anaesthesia. For the DOC dataset having a between-subject design, two-sample t-test (DOC vs. control) was conducted at the group level. The group-level inference was corrected for multiple comparisons using random field theory [voxel level threshold of p<0.005 (uncorrected) and cluster level p<0.05 (FWE-corrected for multiple comparisons)].^67^

Radar plots are additionally presented to indicate the intrinsic connectivity network (ICN) spatial involvement (ICNi) in the contrasts of deep sedation vs. awake and DOC vs. control i.e. shows the voxel overlap between the FC results and canonical ICNs. These canonical ICNs were defined by an atlas^68^ containing 10 well-matched resting-state networks (RSN) from the *ICN_atlas* toolbox (https://www.nitrc.org/projects/icn_atlas/).^69^ The ICNi toolbox calculates the spatial overlap between a connectivity result with the canonical ICNs. Description of ICN-RSN atlas: RSN 1, 2, and 3 (“*visual*”): medial, occipital pole, and lateral visual areas; RSN 4 (“*default mode network*”): medial parietal (precuneus and posterior cingulate), bilateral inferior–lateral–parietal, and ventromedial frontal cortex; RSN 5 (“*cerebellum*”): cerebellum; RSN 6 (“*sensorimotor*”): supplementary motor area, sensorimotor cortex, and secondary somatosensory cortex; RSN 7 (“*auditory*”): superior temporal gyrus, Heschl’s gyrus, and posterior insular. It includes primary and association auditory cortices; RSN 8 (“*executive control*”): medial–frontal areas, including anterior cingulate and paracingulate; RSN 9, 10 (“*frontoparietal*”): frontoparietal areas; these are the only maps to be strongly lateralised. In addition, RSN 9 corresponds strongly to perception–somesthesis–pain, and RSN 10 to cognition–language paradigms, consistent with Broca’s and Wernicke’s areas.^68^ Furthermore, to classify the patterns of thalamo-cortical rs-FC findings based on network involvement (ICNi) with respective to individual nuclei, we performed a hierarchical-based clustering algorithm in RStudio (https://www.r-project.org/).

To investigate which nucleus had the greatest magnitude of change in FC with loss of consciousness, we computed in Matlab the average difference (unsigned magnitude) in FC (ΔFC) across the whole-brain between the baseline (awake condition) and experimental comparison (deep sedation or DOC condition). Mean and standard deviations were used to normalise the values for ΔFC (z-scores), which are displayed in a box plot computed in RStudio, where the y-axis represents the z-scored differential FC and the x-axis depicts the specific thalamic nuclei.

## Results

Our findings address, first and foremost, *which thalamic nuclei* are most strongly associated with loss of consciousness, as well as *how* these changes occur through interactions with cortical networks. We present a whole-brain analysis of thalamo-cortical (and sub-cortical) FC changes across both pharmacological and pathological loss of consciousness. We analysed the fMRI data of N=16 healthy controls under deep sedation (mean plasma propofol concentration of 2.68 𝜇g/mL) and N=22 disorders of consciousness (DOC) patients (N=13 patients in minimally conscious (MCS) state and N=9 in unresponsive wakefulness syndrome (UWS). We used a thalamic parcellation atlas based on DW-MRI, where the thalamus is segmented into 7 regions of interest (ROIs) closely matching the anatomical subparts presented in Morel’s atlas.^41^

### Functional interactions between individual thalamic nuclei and the whole brain in healthy subjects under deep anaesthesia

First, we investigated the spatial localisation of functional connectivity (FC) thalamo-cortical changes with pharmacological (propofol-induced) loss of consciousness by computing whole brain connectivity (normalised as SPM-t) maps following seed-based FC analysis (contrast of deep sedation vs. awake). Visually, SPM-t maps reveal that the Pu and VLV nuclei, but not others, increase their FC with default mode network (DMN) regions, a key hub for consciousness, and decrease their FC with somatomotor (SM)-regions following loss of consciousness (Fig. 1a-i, g-i). By contrast, all other thalamic nuclei exhibited reversed effects with concomitant loss of consciousness (decreased their FC with DMN-regions and increased their connectivity with SM-regions). Correspondingly, the effect was reversed with recovery of consciousness from anaesthesia (Supplementary Fig. 2a-g), validating our findings with DMN and SM-specific patterns of connectivity.

**Fig. 1:**
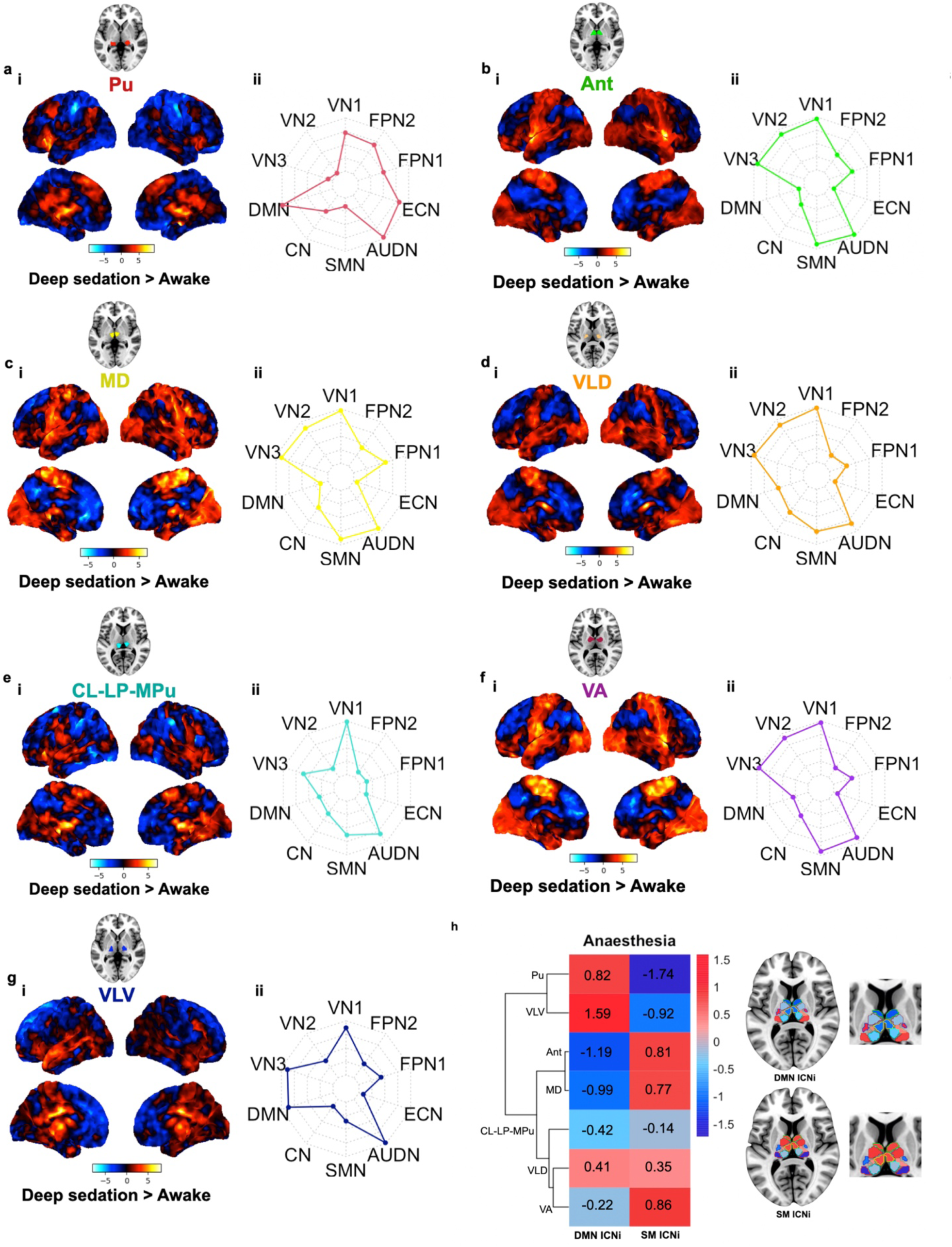
Whole-brain resting-state (rs) functional connectivity (FC) maps in **heathy subjects** in pharmacological (propofol-induced) loss of consciousness (1a-g). Results are computed from seed-to-voxel rs-FC analysis across respective thalamic nuclei [bilateral] seeds — Pu, Ant, MD, VLD, CL-LP-MPu, VA, VLV — and the rest of the brain. Colour bar denotes the strength of the t-statistic. Unthresholded t-maps of the deep sedation vs. awake contrast are shown (**a-i, b-i, c-i, d-i, e-i, f-i, g-i**). Radar plots indicate the intrinsic connectivity network (ICN) spatial involvement (ICNi) of brain regions in the deep sedation vs. awake contrast i.e. shows the voxel overlap between the FC results and canonical ICNs (**a-ii, b-ii, c-ii, d-ii, e-ii, f-ii, g-ii**). These canonical ICNs were defined by an atlas^68^ containing 10 well-matched resting-state networks (RSN) from the *ICN_atlas* toolbox.^69^ Description of ICN-RSN atlas: RSN 1, 2, and 3 (“*visual*”): medial, occipital pole, and lateral visual areas; RSN 4 (“*default mode network*”): medial parietal (precuneus and posterior cingulate), bilateral inferior–lateral–parietal, and ventromedial frontal cortex; RSN 5 (“*cerebellum*”): cerebellum; RSN 6 (“*sensorimotor*”): supplementary motor area, sensorimotor cortex, and secondary somatosensory cortex; RSN 7 (“*auditory*”): superior temporal gyrus, Heschl’s gyrus, and posterior insular. It includes primary and association auditory cortices; RSN 8 (“*executive control*”): medial–frontal areas, including anterior cingulate and paracingulate; RSN 9, 10 (“*frontoparietal*”): frontoparietal areas; these are the only maps to be strongly lateralised. In addition, RSN 9 corresponds strongly to perception–somesthesis–pain, and RSN 10 to cognition–language paradigms, consistent with Broca’s and Wernicke’s areas.^68^ The heatmap (**h**) presents normalised values (z-scores) of ICNi values (DMN and SM from radar plots) for each respective thalamic nucleus following a hierarchical clustering algorithm, which identified the Pu and VLV nuclei into one cluster and all other nuclei into a separate cluster. Concomitant brain maps illustrate ICNi (DMN and SM) for each thalamic nucleus.

Secondly, to quantify more precisely the cortical network involvement in the SPM-t maps, we overlapped our maps with intrinsic brain connectivity networks (ICNs) and calculated the ICN involvement with a resting-state brain network atlas^68^ using the *ICN_atlas* toolbox.^69^ We were able to further validate numerically that, indeed, there was prominent DMN involvement for both the Pu and VLV, exclusively, with pharmacological loss of consciousness (Fig. 1aii-gii). Finally, we used hierarchical clustering to identify these distinctive patterns of functional connectivity we found for Pu and VLV (Fig. 1h). This data-driven result complements our visual and quantitative assessments, as both the Pu and VLV nuclei were grouped into one separate cluster, and all other nuclei were grouped together into a different cluster, providing substantiation to the proposition that, indeed, the Pu and VLV nuclei may manifest more acute contributions than other nuclei in anaesthetic-induced loss of consciousness, through DMN interactions.

### The pulvinar nuclei demonstrated the strongest cortical connectivity change in anaesthetic-induced loss of consciousness

We next sought to investigate which nucleus is most associated with the overall functional connectivity changes in the brain with loss of consciousness. More explicitly, we examined *which* region of the thalamus had the greatest or strongest cortical connectivity change. Thus, to determine which nucleus had the greatest magnitude of change in FC with loss of consciousness, we computed the average change or differential FC (ΔFC) across the whole-brain between the baseline (FC maps indexed by individual’s correlation coefficient *Beta* for ‘awake condition’) and experimental comparison (FC maps for ‘deep sedation condition’). Mean and standard deviations were used to normalise the values for ΔFC (z-scores) displayed in a box plot where the y-axis represents the z-scored ΔFC and the x-axis depicts the specific thalamic nuclei (Fig. 2). Among all thalamic nuclei, the pulvinar (Pu) was found to have the greatest magnitude of ΔFC in healthy subjects with pharmacological loss of consciousness. We then wanted to test whether our preliminary finding that the Pu had the strongest ΔFC — significantly greater change — than the ΔFC of the rest of the nuclei. Thus, a linear mixed model correcting for individual differences was implemented to test for the significance of the Pu connectivity. We compared Pu group against the group of all other nuclei, and indeed, under anaesthesia, the ΔFC for Pu was found to be significant in comparison to the rest of the nuclei (t = 2.081, p = 0.039). [All other nuclei were found to not be significantly different from each other in anaesthesia (p = 0.442) following an F-test (ANOVA)].

**Fig. 2:**
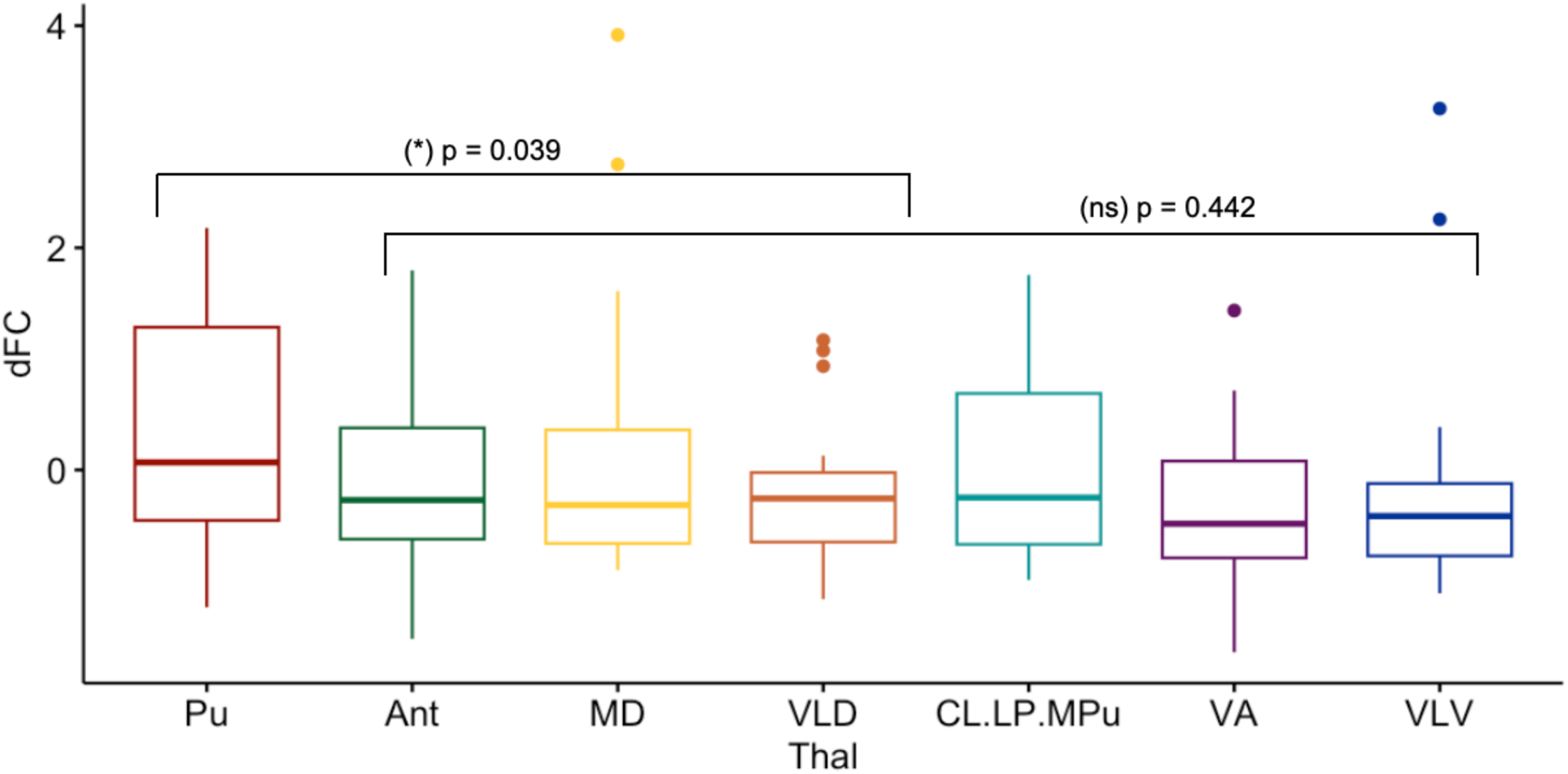
Box plot displays z-scores for the anaesthesia group comparison. Among all nuclei, Pu had the greatest magnitude of ΔFC in anaesthesia. Linear mixed models were used to compare the magnitude of change in FC. Indeed, under anaesthesia, the ΔFC for Pu was found to be statistically significant in comparison to the rest of the nuclei (t = 2.08, p = 0.039*). The rest of the nuclei were found to not be significantly different from each other in anaesthesia (p = 0.442) with an F-test (ANOVA).

Our results on the role of the Pu in overall ΔFC are in line with previous work demonstrating that particularly the medial pulvinar region is involved not only in visuomotor and auditory processing but also associative and higher cognitive processing, and even more so, abnormal medial pulvinar architecture and connectivity were associated to neurodevelopmental disorders.^70^ More relevant to the aims of our research, the medial pulvinar was shown to be involved in altered states of consciousness, notably focal seizures,^71^ and, more recently, stimulation of this region led to improvements in awareness in pharmaco-resistant temporal lobe epilepsy.^72^ Furthermore, several studies demonstrated that lesions to the pulvinar led to changes in conscious content — specifically in impaired feature binding^73^ and hemineglect^74^ — in humans. These insights are further corroborated by our work, as the spatial localisation maps in the previous section (Fig. 1a-g) revealed the Pu has a unique FC profile related to DMN and SM connectivity with loss of consciousness in anaesthesia. What is striking in our dual findings therefore is that the Pu not only has the strongest impact on FC overall in the brain, but that it is strongly coupled with the DMN, a core consciousness hub, suggesting that within the thalamus the Pu nucleus plays a specialised role in pharmacological loss of consciousness.

### Functional interactions between individual thalamic nuclei and the whole brain in DOC patients

We proceeded with our second experimental group to investigate which nucleus in the thalamus (if any) is more closely associated with loss of consciousness in DOC patients. Firstly, we explored the spatial localisation of FC thalamo-cortical changes with pathological loss of consciousness by computing whole brain connectivity (normalised as SPM-t) maps following seed-based FC analysis (contrast of DOC vs. control). Visually, these SPM-t maps revealed the VLV nuclei, but not others, increased their FC with DMN-specific regions (3ai-gi); much like in anaesthesia, the VLV and Pu also decreased their FC with SM-regions (though, in DOC, this was also found with MD, VLD, and CL-LP-MPu). By contrast, all thalamic nuclei other than the VLV exhibited reversed DMN-connectivity profiles with pathologically induced loss of consciousness. Our results are in line with previous resting-state fMRI studies which have demonstrated functional connectivity alterations between DMN areas and the thalamus in brain-injured patients with impaired consciousness.^75,76^ Importantly, previous work has shown significant impairments in the structural connectivity and white matter integrity of the DMN in DOC patients, particularly the pathway connecting posterior cingulate cortex-precuneus to the whole thalamus.^77^

Next, to quantify the cortical network involvement in the SPM-t maps, we overlapped our maps with intrinsic brain connectivity networks (ICNs) and calculated the ICN involvement with a resting-state brain network atlas^68^ using the *ICN_atlas* toolbox.^69^ We were able to further validate numerically that there were substantial functional connectivity gains with the DMN for the VLV, and not other nuclei, with pathological loss of consciousness (Fig. 3aii-gii).

Finally, we used hierarchical clustering to identify these distinctive patterns of functional connectivity we found for Pu and VLV (Fig. 3h). This data-driven result complements our visual and quantitative assessments, as VLV nuclei were grouped into one separate cluster, and all other nuclei were grouped together into a different cluster, providing further evidence to our premise that the VLV, through DMN interactions, may play a more pivotal role in pathological perturbations of consciousness than previously explored. Our findings complement previous lines of evidence highlighting the role of VLV as an integrative centre for motor control, receiving cortical-cerebellar-basal ganglia input and sending core-like projections to primary motor areas.^78,79^

**Fig. 3:**
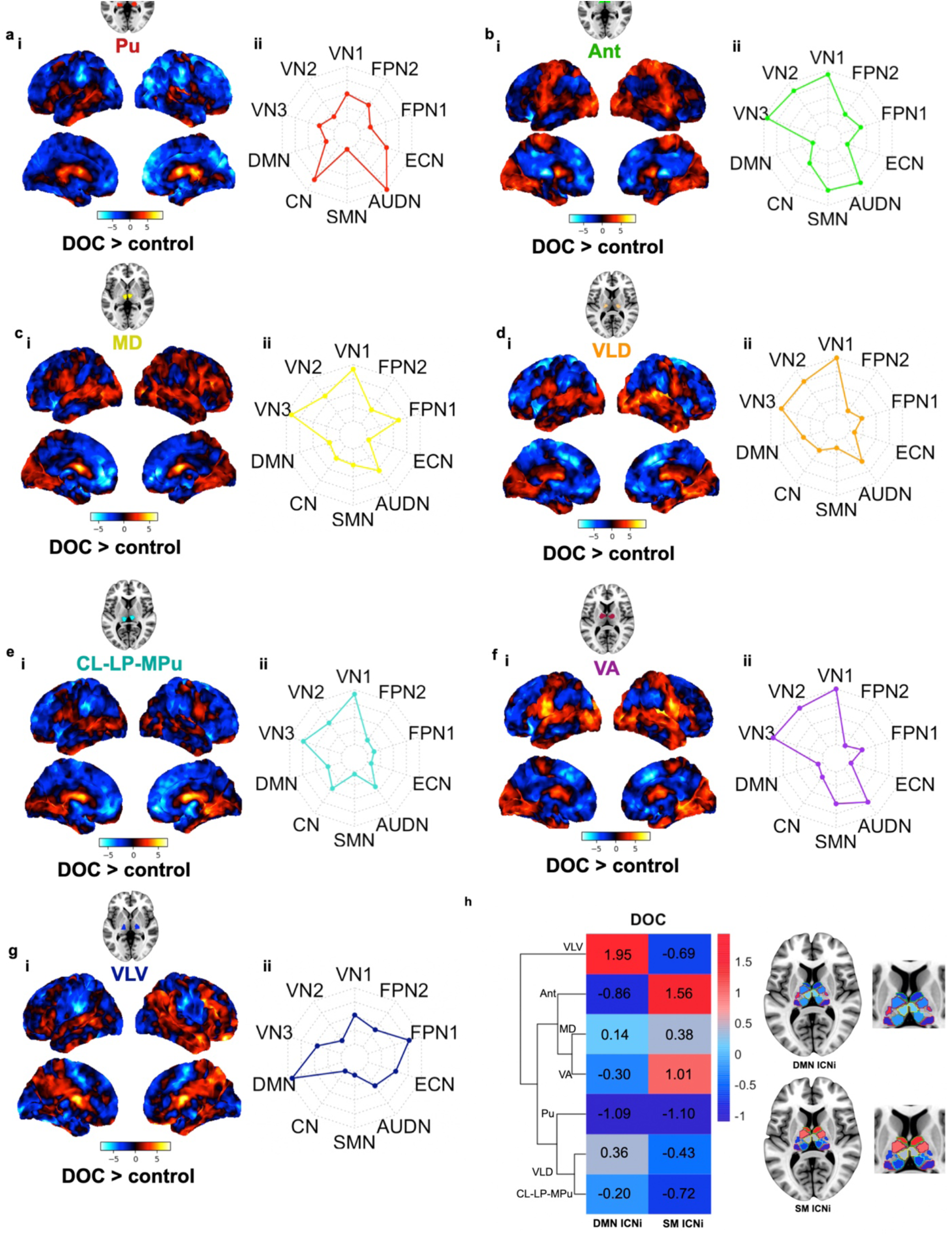
Whole-brain resting-state (rs) functional connectivity (FC) maps in **DOC patients** in loss of consciousness (a-g). Results are computed from seed-to-voxel rs-FC analysis across respective thalamic nuclei [bilateral] seeds — Pu, Ant, MD, VLD, CL-LP-MPu, VA, VLV — and rest of the brain (from a two-sample t-test). Colour bar denotes the strength of the t-statistic. Unthresholded t-maps of the DOC vs. control contrast are shown (**a-i, b-i, c-i, d-i, e-i, f-i, g-i)**. Radar plots indicate the intrinsic connectivity network (ICN) spatial involvement (ICNi) of brain regions in the DOC vs. control contrast i.e. shows the voxel overlap between the FC results and canonical ICNs (**a-ii, b-ii, c-ii, d-ii, e-ii, f-ii, g-ii**). These canonical ICNs were defined by an atlas^68^ containing 10 well-matched resting-state networks (RSN) from the *ICN_atlas* toolbox.^69^ Description of ICN-RSN atlas: RSN 1, 2, and 3 (“*visual*”): medial, occipital pole, and lateral visual areas; RSN 4 (“*default mode network*”): medial parietal (precuneus and posterior cingulate), bilateral inferior–lateral–parietal, and ventromedial frontal cortex; RSN 5 (“*cerebellum*”): cerebellum; RSN 6 (“*sensorimotor*”): supplementary motor area, sensorimotor cortex, and secondary somatosensory cortex; RSN 7 (“*auditory*”): superior temporal gyrus, Heschl’s gyrus, and posterior insular. It includes primary and association auditory cortices; RSN 8 (“*executive control*”): medial–frontal areas, including anterior cingulate and paracingulate; RSN 9, 10 (“*frontoparietal*”): frontoparietal areas; these are the only maps to be strongly lateralised. In addition, RSN 9 corresponds strongly to perception–somesthesis–pain, and RSN 10 to cognition–language paradigms, consistent with Broca’s and Wernicke’s areas.^68^ The heatmap (h) presents normalised values (z-scores) of ICNi values (DMN and SM from radar plots) for each respective thalamic nucleus following a hierarchical clustering algorithm, which identified the VLV nucleus into one cluster and all other nuclei into a separate cluster. Concomitant brain maps illustrate ICNi (DMN and SM) for each thalamic nucleus.

### The ventral-latero-ventral (VLV) nuclei exerted the strongest cortical connectivity change in pathological loss of consciousness in DOC patients

Analogous to our approach in the previous experimental group, we sought to identify which nucleus contributes most to the overall functional connectivity changes in the brain with pathological loss of consciousness. Specifically, we further aimed to distinguish *which* region of the thalamus had the greatest or strongest cortical connectivity change. Therefore, to determine which nucleus had the greatest magnitude of change in FC with loss of consciousness, we computed the average difference in FC (ΔFC) across the whole-brain between the baseline (‘awake condition’) and experimental comparison (‘DOC condition’). Mean and standard deviations were used to normalise the values for ΔFC (z-scores) displayed in a box plot where the y-axis represents the z-scored ΔFC and the x-axis depicts the specific thalamic nuclei (Fig. 4). Among all thalamic nuclei, VLV was found to have the greatest magnitude of ΔFC with pathological loss of consciousness. We then wanted to test whether our preliminary finding that the VLV had the strongest ΔFC is statistically more significant than the ΔFC of the rest of the nuclei. Hence, a linear mixed model correcting for individual differences was implemented to test for the significance of the VLV connectivity. Indeed, in DOC, the ΔFC for VLV was found to be statistically significant in comparison to the rest of the nuclei (t = 12.336, p = 2.2e-16***) with an F-test (ANOVA). Owing to significant differences found among the rest of the nuclei in DOC (p = 1.703e-13), we made multiple comparisons between VLV with each of the rest of the nuclei. After multiple comparison, VLV was significantly the highest among all. We suggest that the generality of significant alterations in the thalamic FC was partly due to the between-subject design of the dataset where the DOC group has considerable individual variations owing to various aetiologies, including TBI, global hypoxic-ischemic encephalopathy, ischaemic stroke, and other causes.^51,80^

**Fig. 4:**
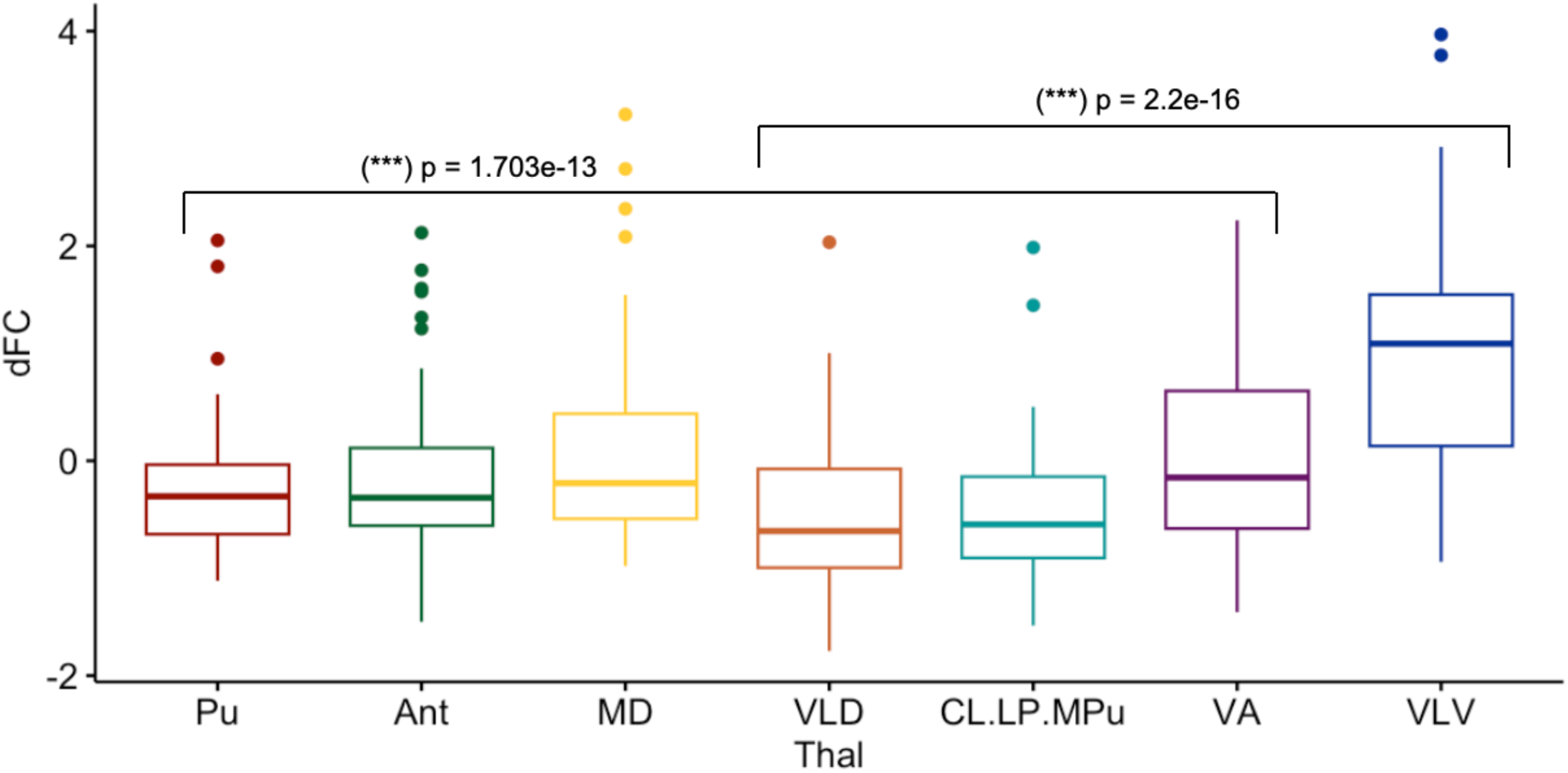
Box plot displays z-scores for the DOC group comparison. Among all nuclei, VLV had the greatest magnitude of ΔFC in DOC. Linear mixed models for within-subject analysis were used to compare the magnitude of change in FC. The ΔFC for VLV was found to be statistically significant in comparison to the rest of the nuclei (t = 12.336, p = 2.2e-16***) with an F-test (ANOVA). Owing to significant differences found among the rest of the nuclei in DOC (p = 1.703e-13), we made multiple comparisons between VLV with each of the rest of the nuclei. After multiple comparison, VLV was significantly the highest among all.

The VLV, known to be part of the motor thalamus, is involved in action control in a circuitry with basal ganglia, motor, premotor, and prefrontal cortices.^81,45,79^ More specifically, VL nuclei have connections to the globus pallidum internal (GPi) of the basal ganglia,^82^ and atrophy in this area has been associated with decreased behavioural arousal patients with disorders of consciousness.^83^ Additionally, a key role for the VL in higher-level cognition, notably in linking attention and memory, has been demonstrated by evidence of VL lesions leading to impairments in stroke patients.^84^ What is striking in our findings thus is that the VLV not only has the strongest impact on FC overall in the brain, but that it is strongly coupled with increases in DMN functional connectivity, a core consciousness hub, suggesting that within the thalamus the VLV nucleus plays a specialised role in pathological loss of consciousness.

In our final analysis (Fig. 5), we sought to further establish thalamic changes in the DOC patients by subjecting them to a test, which assessed their levels of covert consciousness. This test involved tasks of motor imagery (“tennis task”) and spatial imagery (“navigation task”), which allowed us to split the DOC patients into two categories: those with positive responses to the task (“fMRI+ or task responders”) and those with negative responses to the task (“fMRI-or task non-responders”). All positive task-responders (n=7) were TBI patients (except for 1 patient) and were exclusively minimally conscious (MCS) patients. Among those who did not respond to the task (n=15), 6 patients were MCS and 9 patients were in unresponsive wakefulness syndrome (UWS). We used this test as it has been demonstrated to be a more reliable proxy for consciousness because it provides a more sensitive evaluation of the cognitive functions of DOC patients that could not be readily observable with standard behavioural diagnosis such as MCS and UWS.^55,56,85^ This is especially vital given the high rate of misdiagnosis in DOC patients.^86,87^ Concretely, this implies that the subgroup of positive task responders are less severely impaired on the DOC continuum, whereas the subgroup of negative non-responders consists of more severe patients.

**Fig 5:**
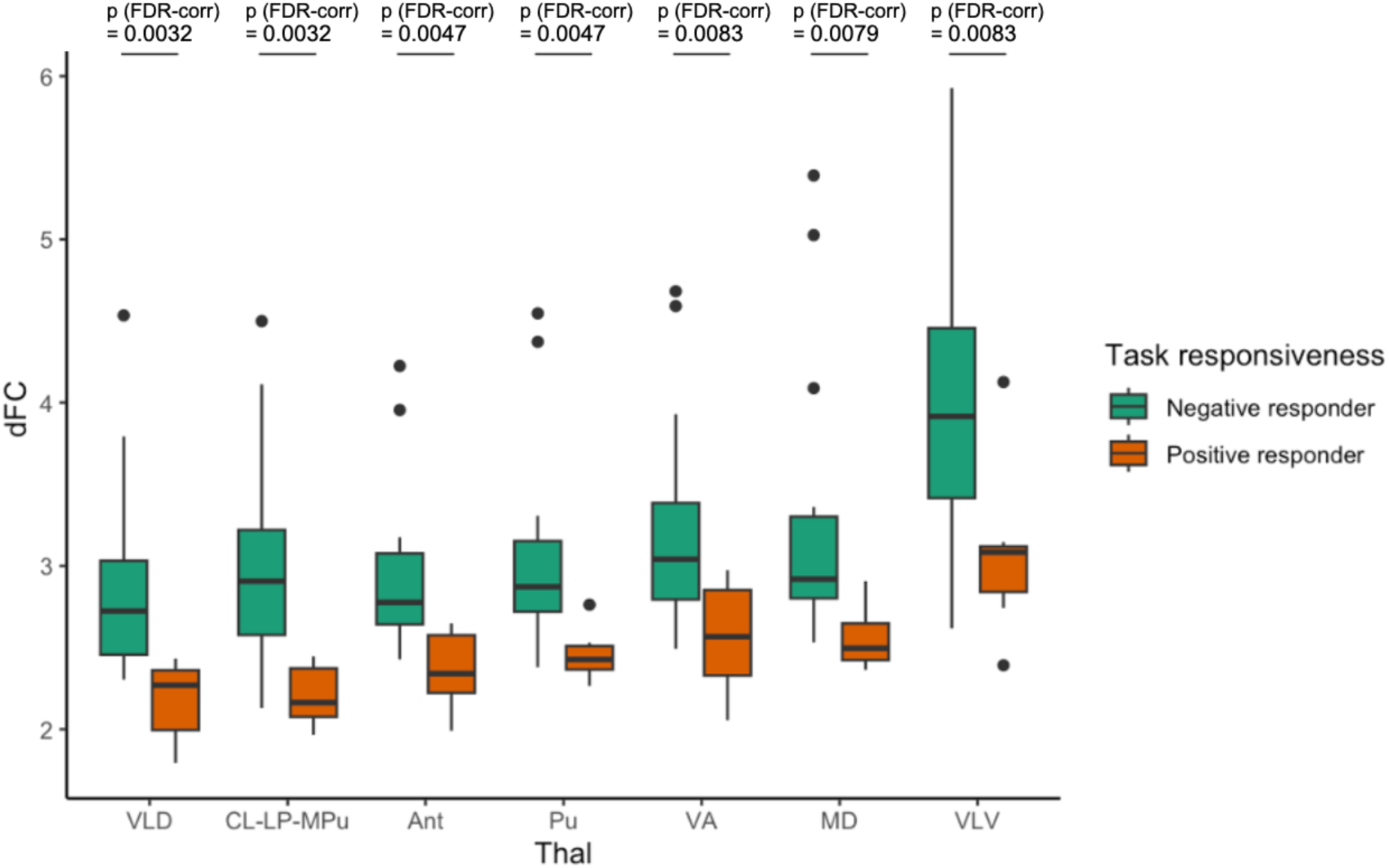
Box plot displays ΔFC scores corresponding to each DOC subgroup (i. fMRI-/task non-responders and ii. fMRI+ /task responders) differential FC from healthy controls. Among all nuclei, the VLV had the greatest ΔFC between task non-responders and task responders after FDR correction.

We proceeded with testing the behavioural relevance of functional connectivity (FC) alterations we found before by investigating the differential functional connectivity (ΔFC) of our thalamic nuclei between each distinct subgroup of DOC patients and healthy controls. The subgroups include those with negative fMRI responses to the Tennis Task (“task non-responders”, fMRI-) and those with positive fMRI responses (“task responders”, fMRI+). Across all seven thalamic nuclei, the ΔFC was found to be statistically significant following a two-sample t-test between the fMRI+ and fMRI-subgroups, corrected for multiple comparisons (FDR-correction). In line with our previous findings (Fig. 4), among all 7 nuclei, the greatest ΔFC across the two subgroups was found for the VLV (p=0.0083) nucleus. Crucially, the VLV is significantly responsible for functional connectivity changes in pathological loss of consciousness overall, and it is even more prominent in the cohort of more severely impaired DOC patients (the negative non-responders). This result reinforces our previous analysis and suggests the utility of VLV, an integral part of the motor thalamus, as not merely a reliable neural marker for consciousness but also, crucially, a behavioural marker that is specific to patients with disorders of consciousness, particularly those are at the more severe end of the spectrum.

## Discussion

We sought to identify the contributions of distinct major thalamic nuclei to perturbed states of consciousness in pharmacological (propofol-induced anaesthesia) and pathological (disorders of consciousness) models in humans. Using fMRI, we examined not only where the largest thalamo-cortical functional connectivity alterations occurred across loss of consciousness but also in which specific thalamic nucleus these changes were most prominent. Whole-brain connectivity maps revealed that, in healthy participants under anaesthesia, the Pu and VLV nuclei increased their functional connectivity with DMN-specific regions and decreased their functional connectivity with SM-regions; all other nuclei exhibited reversed functional connectivity patterns. Furthermore, the alteration of DMN functional connectivity with both Pu and VLV nuclei was reversed in the recovery from the anaesthetised state, which is an important validation of our results. Crucially, among all thalamic nuclei, the Pu was found to have the biggest magnitude of change or strongest differential functional connectivity (ΔFC) with anaesthetic-induced loss of consciousness. In patients with disorders of consciousness, the VLV nuclei increased their functional connectivity with DMN-specific regions and decreased their functional connectivity with SM-regions, with reversed patterns for all other nuclei. Remarkably, in the patient cohort where loss of consciousness was induced by pathological reasons, the VLV was found to have the strongest (ΔFC) among all nuclei. Crucially, we highlight that this finding is not only a prominent neural marker but also reflects changes at the behavioural level, which evidently has important clinical implications.

This research confirms previous literature suggesting that the DMN is a core network that contributes to the contents of consciousness,^15,18^ and provides novel insight into network involvement with subdivisions of the thalamus in both pharmacological and pathological loss of consciousness. Converging evidence has shown the importance of cortical and subcortical contributions of the DMN, and especially posteromedial areas, in healthy subjects under anaesthesia and/or patients with disorders of consciousness.^22,48,80^ Our results align with several resting-state fMRI studies in healthy humans which have demonstrated strong functional connectivity between the precuneus and the whole thalamus during changes in consciousness.^88,89^ Importantly, specific nuclei within the thalamus were found to be structurally and functionally connected to cortical DMN nodes.^89,90,91,92,93^ Furthermore, past work has shown that alterations in structural connectivity in the DMN and along the thalamus-precuneus pathway were correlated with behavioural signs in patients with impaired consciousness,^54^ reinforcing the importance of posteromedial regions in the neural basis of consciousness. Evidence of compromised connectivity between the precuneus and whole-thalamus associated with impaired consciousness have been further demonstrated in various MCS and UWS cohorts.^75,76,77^ Additionally, PET studies also revealed that patients with VS/UWS have altered effective connectivity between the posterior cingulate and frontal association cortices,^94^ as well as between the thalamus, prefrontal and anterior cingulate cortices.^89^ Moreover, network property alterations and functional connectivity disruptions have been found in brain regions associated with consciousness, particularly the medial parietal cortex, frontal cortex, and thalamus in DOC patients.^95^ Furthermore, a functional regulation of the DMN by the thalamus has been suggested in other impaired states of consciousness, specifically during epileptic seizures.^96,97,98^ Clearly, our findings contribute to this growing body of literature and importantly demonstrate that the DMN interacts with specific thalamic nuclei depending on the nature of loss of consciousness, whether pharmacological or pathological.

On a more microscopic level, two distinct cell classes in the thalamus have been shown to be differentially correlated with the cortex, which includes parvalbumin-staining (PVALB) core cells driving excitatory activity via projections to cortical layers III and IV) and calbindin-staining (CALB1) matrix cells projecting diffusely to supra-granular and infragranular layers.^99,100,101^ Intriguingly, previous work showed that CALB1-rich matrix cells in the healthy awake brain exhibited strong preferential functional coupling with several cortical networks, involving the control, limbic, ventral attention, and notably the DMN, whereas in contrast, PVALB-rich core cells favoured functional coupling with visual, dorsal-attention, temporo-parietal, and somatomotor networks.^102^ Moreover, Huang and colleagues observed a shift from a balanced core-matrix unimodal-transmodal functional geometry during consciousness to unimodal core dominance during propofol-induced loss of consciousness in humans.^103^ Additionally, parvalbumin expression was found to be higher in regions with attenuated cortical connectivity during propofol-induced sedation.^104^ There remains a need to further investigate the coupling effects between cortical networks and matrix or core cells in thalamic nuclei in the anaesthetised brain, and even more so, in patients with disorders of consciousness.

Owing to recent advancements in thalamic parcellation,^39,40^ our study has contributed to opening the ‘black box’ of the thalamus and has shown that specific sections of the thalamus are associated with specific types of perturbations of consciousness, pathological or pharmacological. Previously, the thalamus had been explored almost exclusively as a whole entity, and few studies have been able to break down the heterogenous complex nuclei and explore their respective relevance in consciousness in humans.^27,28,29^ In a recent animal study led by Tasserie and colleagues, electrical stimulation of the central thalamus — specifically the central median (CM) nuclei — was found to restore arousal and awareness in a macaque model in propofol-induced loss of consciousness.^25^ Although stimulation of the ventral thalamus control site had no effects on behavioural score in macaques, the authors acknowledged there was no data acquired during an event-related auditory task experiment to measure conscious awareness following ventral lateral thalamic stimulation under anaesthesia^25^, pressing the need for more extended and nuanced investigations. Furthermore, Bastos and colleagues^105^ implanted multiple-contact stimulating electrodes in frontal thalamic nuclei (intralaminar and mediodorsal, with few sites in neighboring ventral posterolateral nucleus) in macaques, and demonstrated that thalamic stimulation reversed the electrophysiologic features of unconsciousness.

Given our findings, not only do we provide a differential picture between pharmacological and pathological states of unconsciousness with relevance to the thalamus, but we also press for the need for more nuanced experimental paradigms that distinguish arousal (alertness or wakefulness) from awareness (content of subjective experience) in human consciousness.^106,107,108^ One model specific to aetiologies of brain injuries producing DOC has been proposed in the clinical literature — the ‘mesocircuit model’ —, which suggests a downregulation of synaptic activity across frontal cortical regions (medial frontal cortices and anterior cingulate cortex), the central thalamus (specifically the central lateral nucleus), and the striatum.^80,109^ Moreover, a functional disfacilitation of the central thalamus, following injuries, reduces activity across projections from this thalamic region to the frontal cortex, posterior medial parietal cortex (a key node of the DMN), and striatum (a subcomponent of the basal ganglia).^5^ Our results provide compelling evidence for a more complex picture in pathological disruptions of consciousness with an important role for the VLV, part of the motor thalamus, possibly through inputs from the globus pallidus and striatum from the basal ganglia circuitry,^79,82,110^ calling for further experimental investigations.

Opening the ‘black box’ of the thalamus significantly contributes to unravelling the ‘black box’ of consciousness. By disentangling the thalamus into more nuanced anatomically-relevant parts, we have been able to suggest that pharmacological and pathological loss of consciousness — although they overlap at the cortical^13,22,26,50,111^ and brainstem^48^ levels — appear to involve distinct thalamic contributions, given that we found the Pu in anaesthesia and VLV in DOC, respectively, to be pivotal. This has clinical implications, as DOC therapies may not be fully reliant upon anaesthetic studies, at least where thalamic connectivity is concerned. Unlike anaesthesia, which taps into the brain connectivity changes through transient neuromodulatory influences, the permanent anatomical damages that accompany brain injury may account for the differential thalamic involvement.^80,112^ By having further separated the DOC in two groups, those severely injured and those less impaired, and by having shown that the VLV remains the predominant nucleus responsible for both, we show consistency within the DOC category of loss of consciousness. Hence, our results not only demonstrate that there is heterogeneity between the consciousness alterations of anaesthesia and DOC in terms of thalamic contributions, but also homogeneity of thalamic involvement within the DOC category, regardless of the exact aetiology of the brain injury and the extent of lesions in the patients. This suggests that, for all its diversity, DOC samples can inform future clinical studies and pinpoint where in the thalamus brain stimulation therapies may prove more efficacious. Given the heterogeneity both in terms of structural damage and functional dysconnectivity in these types of patient cohorts, we would need to, ideally, begin implementing personalised functional connectome mapping^113^ in order to achieve improvements in clinical responses from brain stimulation approaches.^113,114,115^ Our differential findings related to the thalamus depending on the type of unconsciousness is especially pertinent, given recent evidence demonstrating distinctive patterns of cortical engagement between arousable unconsciousness (sleep) and unarousable unconsciousness (propofol-induced anaesthesia),^116^ adding to the debate on the shared versus different mechanisms across multiple conditions of unconsciousness.^117^

Our study has several limitations. Firstly, thalamic subdivisions were not segmented at the basis of individual subjects in their native brain space. The processing pipeline we adopted is standard in neuroimaging studies. The atlas we chose should be able to tolerate the normalization error, as it was shown to have high inter-subject consistency and intra-subject reproducibility.^41^ However, we believe it desirable to generate individual-specific parcellations which could provide even higher anatomic accuracy. This may be experimented and validated by future studies. Secondly, we used the datasets from two imaging sites with non-identical scanner hardware and acquisition parameters. We compared the DOC group to the healthy awake participants from the anaesthesia experiment. However, we are confident this does not influence our interpretation of the main results, due to the fact we focused our analysis on between-nuclei comparisons, by which the system variance caused by different scanners should be evened out. Additionally, our results related to the DOC experiment should hold, because they demonstrated a behavioural correspondence which was tested within subsets of the DOC dataset. In fact, previous publications utilising multi-site datasets have proven the validity of this approach.^22^

Our work which has aimed to identify which thalamic nuclei are more prominent in loss of consciousness could bring diagnostic and therapeutic value for DOC patients. Considering that consciousness depends on the interplay of brain networks,^7,26,118^ we could use localised individualised brain stimulation, in one or multiple nuclei, to induce controlled perturbations in a specific node of the network, which would trigger the propagation of neural activity across proximal and distal brain regions.^119^ In this case, the nuclei which ‘stirred’ the connectivity across the whole brain the most may have the highest chance for neuromodulatory effects following targeted thalamic stimulation. Given that the consensus on optimal therapeutic targets for neuromodulation of consciousness has not yet been established,^80^ several functional neuroimaging studies in patients have been able to suggest the involvement of specific brain regions or circuits for targeted stimulation. One study reverted to cortical stimulation in DOC patients and provided the first proof of principle in a sham-controlled randomized double-blind study that non-invasive transcranial direct current stimulation (tDCS) of anterior cortical regions, notably the dorso-lateral prefrontal cortex (DLPFC), can increase CRS-R scores and lead to cognitive gains in minimally conscious patients (MCS) following brain injury.^120^ Alternatively, central-lateral (CL) thalamic stimulation has been demonstrated in a few single-case studies,^34,121^ and more recently in six patients with disorders of consciousness,^33^ whereas an alternative to DBS, the non-invasive low intensity focused ultrasound (LIFU), targeting the central thalamus has been reported to exhibit clinically significant increases in behavioural responsiveness in two chronic DOC patients.^122,123^ Furthermore, DBS on the central thalamus in a minimally conscious (MCS) patient exhibited reactivations of dormant functional brain networks; however, increases in consciousness were limited.^124^ Moreover, a single-case study demonstrated the effects of DBS were more long-lasting than those observed with pharmacological interventions, notably zolpidem, albeit the behavioural improvements were still limited.^125^ Until now, however, stimulation studies have been limited, which presses for the need for future work – perhaps, we need to be targeting not one but multiple nuclei in the thalamus for recovery of consciousness, as demonstrated by recent investigations of a novel multiple-target thalamic stimulation paradigm in epilepsy.^126^ Given our findings, one way forward could be to redefine stimulation protocols to add the VLV as a target. Our work, thus, offers the most up-to-date elucidation of consciousness-relevant regions in the thalamus, in the hope of marking an accurate, consistent, and reliable target for stimulation site with DBS for recovery from disorders of consciousness in the not-so-distant future.

## Conclusions

In summary, this is a systematic investigation of thalamic connectivity involving both healthy anaesthetised and DOC patients, where we establish that (1) anaesthesia and DOC exhibit distinct involvement of thalamic nuclei for loss of consciousness; (2) different specific nuclei, Pu and VLV, account for the functional interactions in anaesthesia and DOC, respectively. Perhaps most critical of all, we encourage the need for more parcellated explorations into the thalamic mosaic for the purpose of reaching a more unified consensus on the diagnostics and therapies for disorders of consciousness.

## Acknowledgements

This work was supported by grants from the UK Medical Research Council (U.1055.01.002.00001.01) [to A.M.O. and J.D.P.]; The James S. McDonnell Foundation [to A.M.O. and J.D.P.]; The Canada Excellence Research Chairs program (215063) [to A.M.O.]; The Canadian Institute for Advanced Research (CIFAR) [to A.M.O., D.K.M. and E.A.S.]; The National Institute for Health Research (NIHR, UK), Cambridge Biomedical Research Centre and NIHR Senior Investigator Awards [to J.D.P. and D.K.M.]; The British Oxygen Professorship of the Royal College of Anaesthetists [to D.K.M.]; The Evelyn Trust, Cambridge and the EoE CLAHRC fellowship [to J.A.]; The L’Oreal-Unesco for Women in Science Excellence Research Fellowship [to L.N.]; The Stephen Erskine Fellowship, Queens’ College, University of Cambridge [to E.A.S.] and the Gates Cambridge Trust [to A.I.L.]. The research was also supported by the NIHR Brain Injury Healthcare Technology Co-operative based at Cambridge University Hospitals NHS Foundation Trust and University of Cambridge. We would like to thank Victoria Lupson and the staff in the Wolfson Brain Imaging Centre (WBIC) at Addenbrooke’s Hospital for their assistance in scanning.

## Data availability

Due to patient privacy concerns, DOC patient data are available upon request by qualified researchers, for non-commercial use only. The UK Health Research Authority mandates that the confidentiality of data is the responsibility of Chief Investigators for the initial studies (in this case, Dr. Allanson and Prof Menon; and anyone to whom this responsibility is handed – for example, in the context of retirement or transfer to another institution). For researchers interested in working with this dataset, please contact the Data Access Committee: Dr. Judith Allanson, Prof. David Menon or Dr. Emmanuel Stamatakis. The request should include an exact description of the specific planned analysis, the name of the academic institution and department, and the name of the researcher who would bear responsibility. Requests will be considered to assess the feasibility and appropriateness of the proposed study, and the capacity to maintain the required levels of data security, consistent with the original approved Research Ethics approval, and the patient information sheet that was the basis of consent obtained. In case of approval, recipients of the data will be required to a legal agreement confirming that the data will only be used as laid out in the approved proposal, confirming that the data will be protected and not be shared further, and confirming that the data will be deleted as soon as the project finishes. The propofol dataset is available on the OpenNeuro data repository (doi: 10.18112/openneuro.ds003171.v2.0.1).

## Appendix (Supplementary Material)

**Supplementary Fig. 1:**
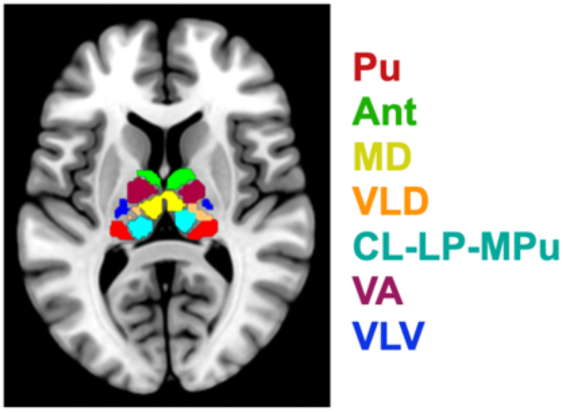
Illustration of 7thalamic masks overlayed onto standard MNI152 template. Thalamic masks correspond to the following bilateral nuclei: pulvinar (Pu), anterior (Ant), medio-dorsal (MD), ventral-latero-dorsal (VLD), central-lateral, lateral-posterior, medial-pulvinar group (CL-LP-MPu), ventral-anterior (VA), and ventral-latero-ventral (VLV) (Najdenovska et al., 2018).

**Supplementary Fig. 2:**
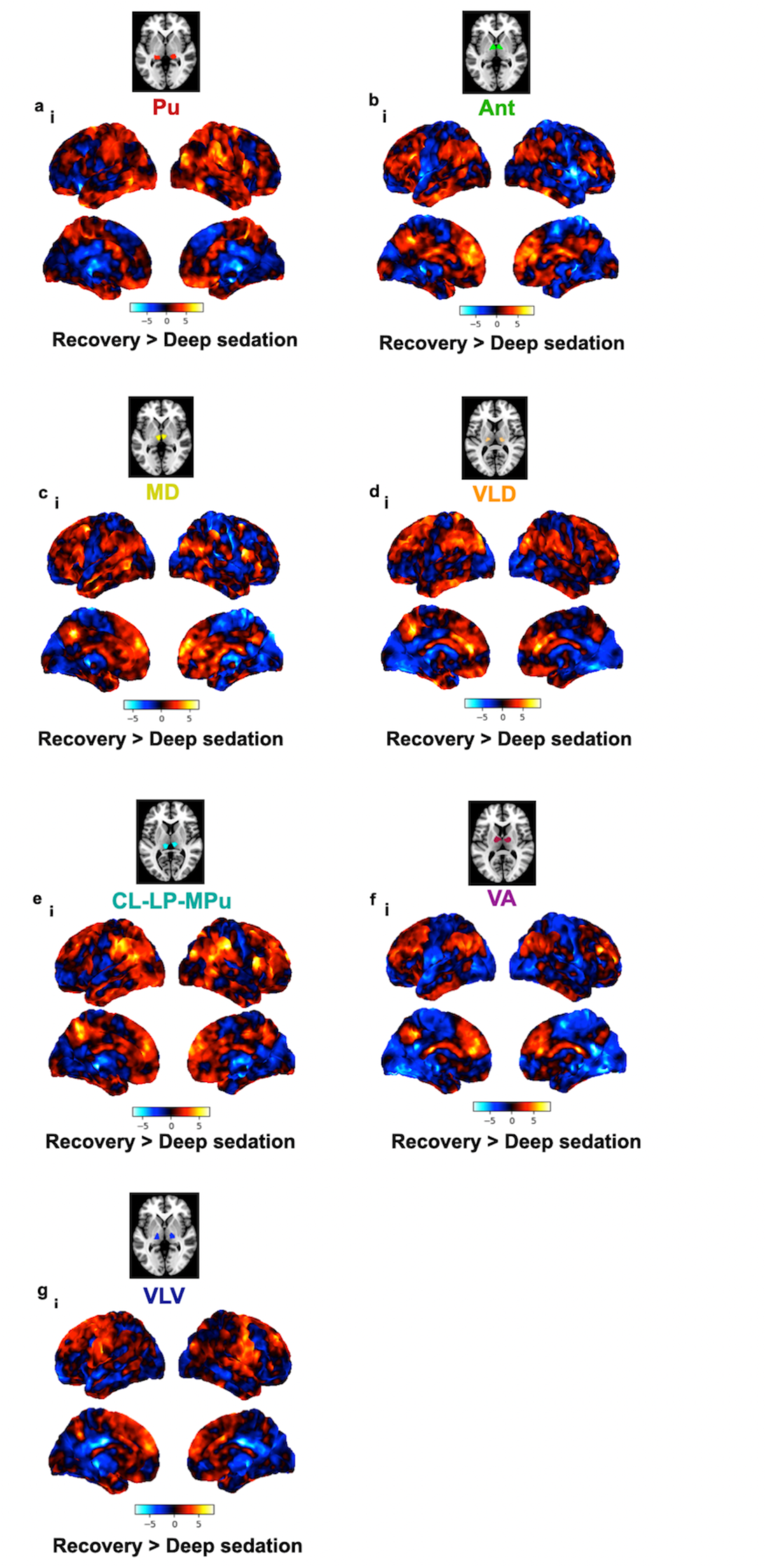
Whole-brain resting-state (rs) functional connectivity (FC) maps in heathy subjects in recovery of consciousness (from pharmacological propofol-induced loss of consciousness). Results are computed from seed-to-voxel rs-FC analysis across respective thalamic nuclei [bilateral] seeds — Pu, Ant, MD, VLD, CL-LP-MPu, VA, VLV — and the rest of the brain. Colour bar denotes the strength of the t-statistic. Unthresholded t-maps of the recovery vs. deep sedation contrast are shown.

## References

1. Avidan, Michael S, and George A Mashour. “Prevention of intraoperative awareness with explicit recall: making sense of the evidence.” Anesthesiology vol. 118,2 (2013): 449–56. doi:10.1097/ALN.0b013e31827ddd2c

2. He, Biyu J. “Next frontiers in consciousness research.” Neuron vol. 111,20 (2023): 3150–3153. doi:10.1016/j.neuron.2023.09.042

3. Owen, Adrian M. “The Search for Consciousness.” Neuron vol. 102,3 (2019): 526–528. doi:10.1016/j.neuron.2019.03.024

4. Shine, James M et al. “The impact of the human thalamus on brain-wide information processing.” Nature reviews. Neuroscience vol. 24,7 (2023): 416–430. doi:10.1038/s41583-023-00701-0

5. Fridman, Esteban A, and Nicholas D Schiff. “Organizing a Rational Approach to Treatments of Disorders of Consciousness Using the Anterior Forebrain Mesocircuit Model.” Journal of clinical neurophysiology : official publication of the American Electroencephalographic Society vol. 39,1 (2022): 40–48. doi:10.1097/WNP.0000000000000729

6. Halassa, Michael M, and S Murray Sherman. “Thalamocortical Circuit Motifs: A General Framework.” Neuron vol. 103,5 (2019): 762–770. doi:10.1016/j.neuron.2019.06.005

7. Hwang, Kai et al. “The Human Thalamus Is an Integrative Hub for Functional Brain Networks.” The Journal of neuroscience : the official journal of the Society for Neuroscience vol. 37,23 (2017): 5594–5607. doi:10.1523/JNEUROSCI.0067-17.2017

8. Shine, James M et al. “The Dynamics of Functional Brain Networks: Integrated Network States during Cognitive Task Performance.” Neuron vol. 92,2 (2016): 544–554. doi:10.1016/j.neuron.2016.09.018

9. Wimmer, Ralf D et al. “Thalamic control of sensory selection in divided attention.” Nature vol. 526,7575 (2015): 705–9. doi:10.1038/nature15398

10. Saalmann, Yuri B et al. “The pulvinar regulates information transmission between cortical areas based on attention demands.” *Science (New York*, N.Y.) vol. 337,6095 (2012): 753–6. doi:10.1126/science.1223082

11. Redinbaugh, Michelle J et al. “Thalamus Modulates Consciousness via Layer-Specific Control of Cortex.” Neuron vol. 106,1 (2020): 66–75.e12. doi:10.1016/j.neuron.2020.01.005

12. Aru, Jaan et al. “Cellular Mechanisms of Conscious Processing.” Trends in cognitive sciences vol. 24,10 (2020): 814–825. doi:10.1016/j.tics.2020.07.006

13. Huang, Zirui et al. “Functional geometry of the cortex encodes dimensions of consciousness.” Nature communications vol. 14,1 72. 5 Jan. 2023, doi:10.1038/s41467-022-35764-7

14. Menon, Vinod. “20 years of the default mode network: A review and synthesis.” Neuron vol. 111,16 (2023): 2469–2487. doi:10.1016/j.neuron.2023.04.023

15. Bodien YG, Threlkeld ZD, Edlow BL. Default mode network dynamics in covert consciousness. Cortex. 2019 Oct;119:571–574. doi: 10.1016/j.cortex.2019.01.014. Epub 2019 Jan 30. PMID: 30791975; PMCID: PMC6527357.

16. Demertzi, Athena et al. “Consciousness supporting networks.” Current opinion in neurobiology vol. 23,2 (2013): 239–44. doi:10.1016/j.conb.2012.12.003

17. Lyu, Dian et al. “A Precuneal Causal Loop Mediates External and Internal Information Integration in the Human Brain.” The Journal of neuroscience : the official journal of the Society for Neuroscience vol. 41,48 (2021): 9944–9956. doi:10.1523/JNEUROSCI.0647-21.2021

18. Yeshurun, Yaara et al. “The default mode network: where the idiosyncratic self meets the shared social world.” Nature reviews. Neuroscience vol. 22,3 (2021): 181–192. doi:10.1038/s41583-020-00420-w

19. Lyu, Dian et al. “Causal evidence for the processing of bodily self in the anterior precuneus.” Neuron vol. 111,16 (2023): 2502–2512.e4. doi:10.1016/j.neuron.2023.05.013

20. Parvizi, Josef et al. “Altered sense of self during seizures in the posteromedial cortex.” Proceedings of the National Academy of Sciences of the United States of America vol. 118,29 (2021): e2100522118. doi:10.1073/pnas.2100522118

21. Herbet, Guillaume et al. “Disrupting posterior cingulate connectivity disconnects consciousness from the external environment.” Neuropsychologia vol. 56 (2014): 239–44. doi:10.1016/j.neuropsychologia.2014.01.020

22. Luppi, Andrea I et al. “Consciousness-specific dynamic interactions of brain integration and functional diversity.” Nature communications vol. 10,1 4616. 10 Oct. 2019, doi:10.1038/s41467-019-12658-9

23. Cui, Yue et al. “Subdivisions of the posteromedial cortex in disorders of consciousness.” NeuroImage. Clinical vol. 20 260–266. 26 Jul. 2018, doi:10.1016/j.nicl.2018.07.025

24. Silva, Stein et al. “Disruption of posteromedial large-scale neural communication predicts recovery from coma.” Neurology vol. 85,23 (2015): 2036–44. doi:10.1212/WNL.0000000000002196

25. Tasserie, Jordy et al. “Deep brain stimulation of the thalamus restores signatures of consciousness in a nonhuman primate model.” Science advances vol. 8,11 (2022): eabl5547. doi:10.1126/sciadv.abl5547

26. Demertzi, A et al. “Human consciousness is supported by dynamic complex patterns of brain signal coordination.” Science advances vol. 5,2 eaat7603. 6 Feb. 2019, doi:10.1126/sciadv.aat7603

27. Weiner, Veronica S et al. “Propofol disrupts alpha dynamics in functionally distinct thalamocortical networks during loss of consciousness.” Proceedings of the National Academy of Sciences of the United States of America vol. 120,11 (2023): e2207831120. doi:10.1073/pnas.2207831120

28. Setzer, Beverly et al. “A temporal sequence of thalamic activity unfolds at transitions in behavioral arousal state.” Nature communications vol. 13,1 5442. 16 Sep. 2022, doi:10.1038/s41467-022-33010-8

29. Liu, Xiaolin et al. “Differential effects of deep sedation with propofol on the specific and nonspecific thalamocortical systems: a functional magnetic resonance imaging study.” Anesthesiology vol. 118,1 (2013): 59–69. doi:10.1097/ALN.0b013e318277a801

30. Graham, D I et al. “Neuropathology of the vegetative state after head injury.” Neuropsychological rehabilitation vol. 15,3–4 (2005): 198–213. doi:10.1080/09602010443000452

31. Maxwell, William L et al. “Thalamic nuclei after human blunt head injury.” Journal of neuropathology and experimental neurology vol. 65,5 (2006): 478–88. doi:10.1097/01.jnen.0000229241.28619.75

32. Whyte, Christopher J et al. “Thalamic contributions to the state and contents of consciousness.” Neuron vol. 112,10 (2024): 1611–1625. doi:10.1016/j.neuron.2024.04.019

33. Schiff, Nicholas D et al. “Thalamic deep brain stimulation in traumatic brain injury: a phase 1, randomized feasibility study.” Nature medicine vol. 29,12 (2023): 3162–3174. doi:10.1038/s41591-023-02638-4

34. Schiff, N D et al. “Behavioural improvements with thalamic stimulation after severe traumatic brain injury.” Nature vol. 448,7153 (2007): 600–3. doi:10.1038/nature06041

35. Karas, Patrick J et al. “Deep Brain Stimulation for Obsessive Compulsive Disorder: Evolution of Surgical Stimulation Target Parallels Changing Model of Dysfunctional Brain Circuits.” Frontiers in neuroscience vol. 12 998. 8 Jan. 2019, doi:10.3389/fnins.2018.00998

36. Cury, Rubens Gisbert et al. “Thalamic deep brain stimulation for tremor in Parkinson disease, essential tremor, and dystonia.” Neurology vol. 89,13 (2017): 1416–1423. doi:10.1212/WNL.0000000000004295

37. Soulier, Hugo et al. “The anterior and pulvinar thalamic nuclei interactions in mesial temporal lobe seizure networks.” Clinical neurophysiology : official journal of the International Federation of Clinical Neurophysiology vol. 150 (2023): 176–183. doi:10.1016/j.clinph.2023.03.016

38. Wu, Teresa Q et al. “Multisite thalamic recordings to characterize seizure propagation in the human brain.” Brain : a journal of neurology vol. 146,7 (2023): 2792–2802. doi:10.1093/brain/awad121

39. Saranathan, Manojkumar et al. “In vivo high-resolution structural MRI-based atlas of human thalamic nuclei.” Scientific data vol. 8,1 275. 28 Oct. 2021, doi:10.1038/s41597-021-01062-y

40. Mai, Jürgen K, and Milan Majtanik. “Toward a Common Terminology for the Thalamus.” Frontiers in neuroanatomy vol. 12 114. 11 Jan. 2019, doi:10.3389/fnana.2018.00114

41. Najdenovska, Elena et al. “In-vivo probabilistic atlas of human thalamic nuclei based on diffusion-weighted magnetic resonance imaging.” Scientific data vol. 5 180270. 27 Nov. 2018, doi:10.1038/sdata.2018.270

42. Iglesias, Juan Eugenio et al. “A probabilistic atlas of the human thalamic nuclei combining ex vivo MRI and histology.” NeuroImage vol. 183 (2018): 314–326. doi:10.1016/j.neuroimage.2018.08.012

43. Kumar, Vinod et al. “Direct diffusion-based parcellation of the human thalamus.” Brain structure & function vol. 220,3 (2015): 1619–35. doi:10.1007/s00429-014-0748-2

44. Zhang, Dongyang et al. “Intrinsic functional relations between human cerebral cortex and thalamus.” Journal of neurophysiology vol. 100,4 (2008): 1740–8. doi:10.1152/jn.90463.2008

45. Behrens, T E J et al. “Non-invasive mapping of connections between human thalamus and cortex using diffusion imaging.” Nature neuroscience vol. 6,7 (2003): 750–7. doi:10.1038/nn1075

46. Luppi, Andrea I et al. “Distributed harmonic patterns of structure-function dependence orchestrate human consciousness.” Communications biology vol. 6,1 117. 28 Jan. 2023, doi:10.1038/s42003-023-04474-1

47. Coppola, Peter et al. “Network dynamics scale with levels of awareness.” NeuroImage vol. 254 (2022): 119128. doi:10.1016/j.neuroimage.2022.119128

48. Spindler, Lennart R B et al. “Dopaminergic brainstem disconnection is common to pharmacological and pathological consciousness perturbation.” Proceedings of the National Academy of Sciences of the United States of America vol. 118,30 (2021): e2026289118. doi:10.1073/pnas.2026289118

49. Naci, Lorina et al. “Functional diversity of brain networks supports consciousness and verbal intelligence.” Scientific reports vol. 8,1 13259. 5 Sep. 2018, doi:10.1038/s41598-018-31525-z

50. Luppi, Andrea I et al. “Whole-brain modelling identifies distinct but convergent paths to unconsciousness in anaesthesia and disorders of consciousness.” Communications biology vol. 5,1 384. 20 Apr. 2022, doi:10.1038/s42003-022-03330-y

51. Edlow, Brian L et al. “Early detection of consciousness in patients with acute severe traumatic brain injury.” Brain : a journal of neurology vol. 140,9 (2017): 2399–2414. doi:10.1093/brain/awx176

52. Varley, T. F., Luppi, A. I., Pappas, I., Naci, L., Adapa, R., Owen, A. M., Menon, D. K., & Stamatakis, E. A. (2020). Consciousness & Brain Functional Complexity in Propofol Anaesthesia. Scientific reports, 10(1), 1018. 10.1038/s41598-020-57695-3

53. Craig, M. M., et al. “Resting-state based prediction of task-related activation in patients with disorders of consciousness.” Preprint at bioRxiv (2021): 10.1101/2021.03.27.436534

54. Fernández-Espejo, Davinia et al. “The clinical utility of fMRI for identifying covert awareness in the vegetative state: a comparison of sensitivity between 3T and 1.5T.” PloS one vol. 9,4 e95082. 14 Apr. 2014, doi:10.1371/journal.pone.0095082

55. Monti, Martin M et al. “Willful modulation of brain activity in disorders of consciousness.” The New England journal of medicine vol. 362,7 (2010): 579–89. doi:10.1056/NEJMoa0905370

56. Owen, Adrian M et al. “Detecting awareness in the vegetative state.” *Science (New York*, N.Y.) vol. 313,5792 (2006): 1402. doi:10.1126/science.1130197

57. Calhoun, Vince D et al. “The impact of T1 versus EPI spatial normalization templates for fMRI data analyses.” Human brain mapping vol. 38,11 (2017): 5331–5342. doi:10.1002/hbm.23737

58. Ashburner, John, and Karl J Friston. “Unified segmentation.” NeuroImage vol. 26,3 (2005): 839–51. doi:10.1016/j.neuroimage.2005.02.018

59. Power, Jonathan D et al. “Ridding fMRI data of motion-related influences: Removal of signals with distinct spatial and physical bases in multiecho data.” Proceedings of the National Academy of Sciences of the United States of America vol. 115,9 (2018): E2105–E2114. doi:10.1073/pnas.1720985115

60. Weiler, Marina et al. “Evaluating denoising strategies in resting-state functional magnetic resonance in traumatic brain injury (EpiBioS4Rx).” Human brain mapping vol. 43,15 (2022): 4640–4649. doi:10.1002/hbm.25979

61. Crone, Julia S et al. “A systematic investigation of the association between network dynamics in the human brain and the state of consciousness.” Neuroscience of consciousness vol. 2020,1 niaa008. 14 Jun. 2020, doi:10.1093/nc/niaa008

62. Parkes, Linden et al. “An evaluation of the efficacy, reliability, and sensitivity of motion correction strategies for resting-state functional MRI.” NeuroImage vol. 171 (2018): 415–436. doi:10.1016/j.neuroimage.2017.12.073

63. Whitfield-Gabrieli, Susan, and Alfonso Nieto-Castanon. “Conn: a functional connectivity toolbox for correlated and anticorrelated brain networks.” Brain connectivity vol. 2,3 (2012): 125–41. doi:10.1089/brain.2012.0073

64. Battistella, Giovanni et al. “Robust thalamic nuclei segmentation method based on local diffusion magnetic resonance properties.” Brain structure & function vol. 222,5 (2017): 2203–2216. doi:10.1007/s00429-016-1336-4

65. Morel, A et al. “Multiarchitectonic and stereotactic atlas of the human thalamus.” The Journal of comparative neurology vol. 387,4 (1997): 588–630. doi:10.1002/(sici)1096-9861(19971103)387:4<588::aid-cne8>3.0.co;2-z

66. Woodrow, Rebecca E et al. “Acute thalamic connectivity precedes chronic post-concussive symptoms in mild traumatic brain injury.” Brain : a journal of neurology vol. 146,8 (2023): 3484–3499. doi:10.1093/brain/awad056

67. Eklund, Anders et al. “Cluster failure: Why fMRI inferences for spatial extent have inflated false-positive rates.” Proceedings of the National Academy of Sciences of the United States of America vol. 113,28 (2016): 7900–5. doi:10.1073/pnas.1602413113

68. Smith, Stephen M et al. “Correspondence of the brain’s functional architecture during activation and rest.” Proceedings of the National Academy of Sciences of the United States of America vol. 106,31 (2009): 13040–5. doi:10.1073/pnas.0905267106

69. Kozák, Lajos R et al. “ICN_Atlas: Automated description and quantification of functional MRI activation patterns in the framework of intrinsic connectivity networks.” NeuroImage vol. 163 (2017): 319–341. doi:10.1016/j.neuroimage.2017.09.014

70. Homman-Ludiye, Jihane, and James A Bourne. “The medial pulvinar: function, origin and association with neurodevelopmental disorders.” Journal of anatomy vol. 235,3 (2019): 507–520. doi:10.1111/joa.12932

71. Filipescu, Cristina et al. “The effect of medial pulvinar stimulation on temporal lobe seizures.” Epilepsia vol. 60,4 (2019): e25–e30. doi:10.1111/epi.14677

72. Deutschová, Barbora et al. “Ictal connectivity changes induced by pulvinar stimulation correlate with improvement of awareness.” Brain stimulation vol. 14,2 (2021): 344–346. doi:10.1016/j.brs.2021.01.021

73. Ward, Robert et al. “Deficits in spatial coding and feature binding following damage to spatiotopic maps in the human pulvinar.” Nature neuroscience vol. 5,2 (2002): 99–100. doi:10.1038/nn794

74. Karnath, Hans Otto et al. “The subcortical anatomy of human spatial neglect: putamen, caudate nucleus and pulvinar.” Brain : a journal of neurology vol. 125,Pt 2 (2002): 350–60. doi:10.1093/brain/awf032

75. Crone, Julia Sophia et al. “Altered network properties of the fronto-parietal network and the thalamus in impaired consciousness.” NeuroImage. Clinical vol. 4 240–8. 26 Dec. 2013, doi:10.1016/j.nicl.2013.12.005

76. Vanhaudenhuyse, Audrey et al. “Default network connectivity reflects the level of consciousness in non-communicative brain-damaged patients.” Brain : a journal of neurology vol. 133,Pt 1 (2010): 161–71. doi:10.1093/brain/awp313

77. Fernández-Espejo, Davinia et al. “A role for the default mode network in the bases of disorders of consciousness.” Annals of neurology vol. 72,3 (2012): 335–43. doi:10.1002/ana.23635

78. Shepherd, Gordon M G, and Naoki Yamawaki. “Untangling the cortico-thalamo-cortical loop: cellular pieces of a knotty circuit puzzle.” Nature reviews. Neuroscience vol. 22,7 (2021): 389–406. doi:10.1038/s41583-021-00459-3

79. McFarland, Nikolaus R, and Suzanne N Haber. “Thalamic relay nuclei of the basal ganglia form both reciprocal and nonreciprocal cortical connections, linking multiple frontal cortical areas.” The Journal of neuroscience : the official journal of the Society for Neuroscience vol. 22,18 (2002): 8117–32. doi:10.1523/JNEUROSCI.22-18-08117.2002

80. Edlow, Brian L et al. “Recovery from disorders of consciousness: mechanisms, prognosis and emerging therapies.” Nature reviews. Neurology vol. 17,3 (2021): 135–156. doi:10.1038/s41582-020-00428-x

81. Wolff, Mathieu, and Seralynne D Vann. “The Cognitive Thalamus as a Gateway to Mental Representations.” The Journal of neuroscience : the official journal of the Society for Neuroscience vol. 39,1 (2019): 3–14. doi:10.1523/JNEUROSCI.0479-18.2018

82. Zheng, Zhong S, and Martin M Monti. “Cortical and thalamic connections of the human globus pallidus: Implications for disorders of consciousness.” Frontiers in neuroanatomy vol. 16 960439. 25 Aug. 2022, doi:10.3389/fnana.2022.960439

83. Lutkenhoff, Evan S et al. “Thalamic and extrathalamic mechanisms of consciousness after severe brain injury.” Annals of neurology vol. 78,1 (2015): 68–76. doi:10.1002/ana.24423

84. de Bourbon-Teles, José et al. “Thalamic control of human attention driven by memory and learning.” Current biology : CB vol. 24,9 (2014): 993–9. doi:10.1016/j.cub.2014.03.024

85. Schnakers, Caroline et al. “Diagnostic accuracy of the vegetative and minimally conscious state: clinical consensus versus standardized neurobehavioral assessment.” BMC neurology vol. 9 35. 21 Jul. 2009, doi:10.1186/1471-2377-9-35

86. Wang, Jing et al. “The misdiagnosis of prolonged disorders of consciousness by a clinical consensus compared with repeated coma-recovery scale-revised assessment.” BMC neurology vol. 20,1 343. 12 Sep. 2020, doi:10.1186/s12883-020-01924-9

87. Peterson, Andrew et al. “Risk, diagnostic error, and the clinical science of consciousness.” NeuroImage. Clinical vol. 7 588–97. 20 Feb. 2015, doi:10.1016/j.nicl.2015.02.008

88. Zhang, Sheng, and Chiang-shan R Li. “Functional connectivity mapping of the human precuneus by resting state fMRI.” NeuroImage vol. 59,4 (2012): 3548–62. doi:10.1016/j.neuroimage.2011.11.023

89. Cunningham, Samantha I et al. “Structural and functional connectivity of the precuneus and thalamus to the default mode network.” Human brain mapping vol. 38,2 (2017): 938–956. doi:10.1002/hbm.23429

90. Alves, Pedro Nascimento et al. “An improved neuroanatomical model of the default-mode network reconciles previous neuroimaging and neuropathological findings.” Communications biology vol. 2 370. 10 Oct. 2019, doi:10.1038/s42003-019-0611-3

91. Lee, Tien-Wen, and Shao-Wei Xue. “Functional connectivity maps based on hippocampal and thalamic dynamics may account for the default-mode network.” The European journal of neuroscience vol. 47,5 (2018): 388–398. doi:10.1111/ejn.13828

92. Parvizi, Josef et al. “Neural connections of the posteromedial cortex in the macaque.” Proceedings of the National Academy of Sciences of the United States of America vol. 103,5 (2006): 1563–8. doi:10.1073/pnas.0507729103

93. Cauda, Franco et al. “Functional connectivity of the posteromedial cortex.” PloS one vol. 5,9 e13107. 30 Sep. 2010, doi:10.1371/journal.pone.0013107

94. Laureys, S et al. “Impaired effective cortical connectivity in vegetative state: preliminary investigation using PET.” NeuroImage vol. 9,4 (1999): 377–82. doi:10.1006/nimg.1998.0414

95. Laureys, S et al. “Restoration of thalamocortical connectivity after recovery from persistent vegetative state.” *Lancet (London*, England) vol. 355,9217 (2000): 1790–1. doi:10.1016/s0140-6736(00)02271-6

96. Herbet, Guillaume et al. “Disrupting posterior cingulate connectivity disconnects consciousness from the external environment.” Neuropsychologia vol. 56 (2014): 239-44. doi:10.1016/j.neuropsychologia.2014.01.020

97. Danielson, Nathan B et al. “The default mode network and altered consciousness in epilepsy.” Behavioural neurology vol. 24,1 (2011): 55–65. doi:10.3233/BEN-2011-0310

98. Blumenfeld, H et al. “Cortical and subcortical networks in human secondarily generalized tonic-clonic seizures.” Brain : a journal of neurology vol. 132,Pt 4 (2009): 999–1012. doi:10.1093/brain/awp028

99. Müller, Eli J et al. “Core and matrix thalamic sub-populations relate to spatio-temporal cortical connectivity gradients.” NeuroImage vol. 222 (2020): 117224. doi:10.1016/j.neuroimage.2020.117224

100. Clascá, Francisco et al. “Unveiling the diversity of thalamocortical neuron subtypes.” The European journal of neuroscience vol. 35,10 (2012): 1524–32. doi:10.1111/j.1460-9568.2012.08033.x

101. Jones, E G. “The thalamic matrix and thalamocortical synchrony.” Trends in neurosciences vol. 24,10 (2001): 595–601. doi:10.1016/s0166-2236(00)01922-6

102. Müller, Eli J et al. “The non-specific matrix thalamus facilitates the cortical information processing modes relevant for conscious awareness.” Cell reports vol. 42,8 (2023): 112844. doi:10.1016/j.celrep.2023.112844

103. Huang, Zirui, et al. “Propofol Disrupts the Functional Core-Matrix Architecture of the Thalamus in Humans.” *bioRxiv :* the preprint server for biology 2024.01.23.576934. 24 Jan. 2024, doi:10.1101/2024.01.23.576934. Preprint.

104. Craig, Michael M et al. “Propofol sedation-induced alterations in brain connectivity reflect parvalbumin interneurone distribution in human cerebral cortex.” British journal of anaesthesia vol. 126,4 (2021): 835–844. doi:10.1016/j.bja.2020.11.035

105. Bastos, André M et al. “Neural effects of propofol-induced unconsciousness and its reversal using thalamic stimulation.” eLife vol. 10 e60824. 27 Apr. 2021, doi:10.7554/eLife.60824

106. Lee, Minji et al. “Quantifying arousal and awareness in altered states of consciousness using interpretable deep learning.” Nature communications vol. 13,1 1064. 25 Feb. 2022, doi:10.1038/s41467-022-28451-0

107. Koch, Christof et al. “Neural correlates of consciousness: progress and problems.” Nature reviews. Neuroscience vol. 17,5 (2016): 307–21. doi:10.1038/nrn.2016.22

108. Tononi, Giulio et al. “Integrated information theory: from consciousness to its physical substrate.” Nature reviews. Neuroscience vol. 17,7 (2016): 450–61. doi:10.1038/nrn.2016.44

109. Schiff, Nicholas D. “Recovery of consciousness after brain injury: a mesocircuit hypothesis.” Trends in neurosciences vol. 33,1 (2010): 1–9. doi:10.1016/j.tins.2009.11.002

110. Shine, James M. “The thalamus integrates the macrosystems of the brain to facilitate complex, adaptive brain network dynamics.” Progress in neurobiology vol. 199 (2021): 101951. doi:10.1016/j.pneurobio.2020.101951

111. Huang, Zirui et al. “Temporal circuit of macroscale dynamic brain activity supports human consciousness.” Science advances vol. 6,11 eaaz0087. 11 Mar. 2020, doi:10.1126/sciadv.aaz0087

112. Raguž, Marina et al. “Structural changes in brains of patients with disorders of consciousness treated with deep brain stimulation.” Scientific reports vol. 11,14401. 23 Feb. 2021, doi:10.1038/s41598-021-83873-y

113. Edlow, Brian L et al. “Personalized Connectome Mapping to Guide Targeted Therapy and Promote Recovery of Consciousness in the Intensive Care Unit.” Neurocritical care vol. 33,2 (2020): 364–375. doi:10.1007/s12028-020-01062-7

114. Provencio, J Javier, et al. “The Curing Coma Campaign: Framing Initial Scientific Challenges-Proceedings of the First Curing Coma Campaign Scientific Advisory Council Meeting.” Neurocritical care vol. 33,1 (2020): 1–12. doi:10.1007/s12028-020-01028-9

115. Thibaut, Aurore et al. “Therapeutic interventions in patients with prolonged disorders of consciousness.” The Lancet. Neurology vol. 18,6 (2019): 600–614. doi:10.1016/S1474-4422(19)30031-6

116. Zelmann, Rina et al. “Differential cortical network engagement during states of un/consciousness in humans.” Neuron vol. 111,21 (2023): 3479–3495.e6. doi:10.1016/j.neuron.2023.08.007

117. Eikermann, Matthias et al. “Sleep and Anesthesia: The Shared Circuit Hypothesis Has Been Put to Bed.” Current biology : CB vol. 30,5 (2020): R219–R221. doi:10.1016/j.cub.2020.01.057

118. Li, Hui et al. “Functional networks in prolonged disorders of consciousness.” Frontiers in neuroscience vol. 17 1113695. 17 Feb. 2023, doi:10.3389/fnins.2023.1113695

119. Ozdemir, Recep A et al. “Individualized perturbation of the human connectome reveals reproducible biomarkers of network dynamics relevant to cognition.” Proceedings of the National Academy of Sciences of the United States of America vol. 117,14 (2020): 8115–8125. doi:10.1073/pnas.1911240117

120. Thibaut, Aurore et al. “tDCS in patients with disorders of consciousness: sham-controlled randomized double-blind study.” Neurology vol. 82,13 (2014): 1112–8. doi:10.1212/WNL.0000000000000260

121. Gottshall, Jackie L et al. “Daytime Central Thalamic Deep Brain Stimulation Modulates Sleep Dynamics in the Severely Injured Brain: Mechanistic Insights and a Novel Framework for Alpha-Delta Sleep Generation.” Frontiers in neurology vol. 10 20. 4 Feb. 2019, doi:10.3389/fneur.2019.00020

122. Cain, Josh A et al. “Ultrasonic Deep Brain Neuromodulation in Acute Disorders of Consciousness: A Proof-of-Concept.” Brain sciences vol. 12,4 428. 23 Mar. 2022, doi:10.3390/brainsci12040428

123. Monti, Martin M et al. “Non-Invasive Ultrasonic Thalamic Stimulation in Disorders of Consciousness after Severe Brain Injury: A First-in-Man Report.” Brain stimulation vol. 9,6 (2016): 940–941. doi:10.1016/j.brs.2016.07.008

124. Arnts, Hisse et al. “Clinical and neurophysiological effects of central thalamic deep brain stimulation in the minimally conscious state after severe brain injury.” Scientific reports vol. 12,1 12932. 28 Jul. 2022, doi:10.1038/s41598-022-16470-2

125. Arnts, Hisse et al. “Deep brain stimulation of the central thalamus restores arousal and motivation in a zolpidem-responsive patient with akinetic mutism after severe brain injury.” Scientific reports vol. 14,12950. 5 Feb. 2024, doi:10.1038/s41598-024-52267-1

126. Yang, Andrew I et al. “Multitarget deep brain stimulation for epilepsy.” Journal of neurosurgery vol. 140,1 210–217. 14 Jul. 2023, doi:10.3171/2023.5.JNS23982

